# nAPOGEE: A machine-learning platform for clinically actionable pathogenicity assessment of all mitochondrial noncoding variants

**DOI:** 10.64898/2025.12.20.695671

**Authors:** Salvatore Daniele Bianco, Agnese Giovannetti, Marta Adinolfi, Francesco Petrizzelli, Andrea Villani, Manuel Mangoni, Tommaso Biagini, Niccolò Liorni, Alessandro Napoli, Vincent Procaccio, Marie T. Lott, Shiping Zhang, Leonardo Caporali, Daniele Ghezzi, Andrea Urbani, Antonio Gasbarrini, Douglas C. Wallace, Viviana Caputo, Tommaso Mazza

## Abstract

Mitochondrial noncoding variants, particularly those in tRNA and rRNA genes, pose significant challenges for clinical interpretation due to heteroplasmy, broad phenotypic heterogeneity where symptoms can overlap with other conditions, and the limited availability of well-established genotype-phenotype correlations. Despite their central role in mitochondrial translation, these variants have remained largely unexplored by the existing variant-effect predictors. Here, we present nAPOGEE, a novel machine-learning framework specifically designed to assess the pathogenicity of all possible single-nucleotide variants in human mitochondrial noncoding RNAs. nAPOGEE integrates two specialized predictors: tAPOGEE, which outperforms existing tools for tRNAs, and rAPOGEE, the first dedicated classifier for mitochondrial rRNA variants. Using curated training datasets, phylogenetic conservation metrics, secondary structure modeling, RNA-specific embeddings, and thermodynamic features, nAPOGEE provides biologically interpretable predictions and posterior probabilities aligned with the ACMG/AMP guidelines. Applied to both curated variant sets and population-scale data, nAPOGEE revealed consistent spatial correlation of the predicted pathogenicity, reflecting underlying structural and evolutionary constraints. This study addresses a longstanding gap in mitochondrial genomics and offers a clinically applicable tool for variant prioritization, reclassification, and research into mitochondrial disease mechanisms.

## Introduction

Redundancy refers to the existence of multiple gene copies that serve similar or identical purposes. This concept is particularly significant for transfer RNA (tRNA) genes in the nuclear genome, where redundancy is essential for maintaining cellular functions and reducing the impact of potentially harmful mutations. Numerous tRNA genes exist in multiple copies and are often distributed across various chromosomal sites. This redundancy, which is argued to follow cell-specific pattern^1^, can manifest as replicated genes, which serve as backups if one copy becomes inoperative owing to mutation, and tRNA gene groups, which code for tRNAs with distinct anticodons but can still recognize identical or similar codons, thus compensating for each other. Redundancy in tRNA molecules serves two purposes: it enables cells to adjust the tRNA supply based on cellular needs and prevents a single tRNA gene mutation from severely disrupting protein synthesis. As a result, no human diseases were found to be associated with mutations in the nuclear-encoded cytosolic tRNAs sequences, except for one atypical case. A homozygous single-nucleotide variant in tRNA for selenocysteine incorporation (tRNA-Sec) was found in a young male with symptoms such as abdominal pain, weakness, fatigue, thyroid dysfunction, and low plasma selenium levels, resembling other mutations affecting selenoprotein synthesis^2^. tRNA-Sec is a single-copy gene in the nuclear genome and its removal from the mouse genome results in embryonic lethality^3^. In contrast, as the various steps involved in tRNA processing, including splicing, modification, maturation, and charging, play a crucial role in determining tRNA function, when defects occur in these processes, they can affect a large number of tRNAs, thereby overcoming the normal functional redundancy and ultimately leading to dysfunction. For example, both biallelic and dominant variants have been found in aminoacyl-tRNA synthetase (aaRS) genes, leading to various neurological and systemic disorders^4^. Many nuclear tRNA-related disorders predominantly affect the nervous system; however, the reasons for this sensitivity are not fully understood. Hence, the apparent absence of disease-causing mutations in typical nuclear tRNA genes aligns with the concept of functional redundancy among isodecoder tRNAs, that is, tRNA molecules that share the same anticodon but differ in their sequence elsewhere in the molecule. Notably, the number of isodecoder tRNA genes decreases in less complex organisms, with yeast exhibiting minimal redundancy^1,5^. Consequently, mutations in nuclear tRNA genes are likely to produce observable phenotypes in simpler organisms^1^.

Similarly, multiple copies of nuclear ribosomal RNA (rRNA) genes have been identified due to their essential roles in protein synthesis. Humans have, in fact, approximately 300–400 copies of rRNA genes in their nuclear genome which are organized in tandem repeats located on the short arms of five acrocentric chromosomes^6^. Not all these genes are actively transcribed simultaneously. Some copies are silent or inactive and are regulated by epigenetic modifications such as DNA methylation and histone modifications^7^. More complex organisms have evolved a less economical form of rRNA synthesis, with plants and lower animals up to reptiles exhibiting a relatively small rRNA transcription unit (2.7-2.8 million Daltons) in contrast with birds and mammals that have a larger transcription unit (4.0-4.2 million Daltons)^8^. Although this potentially allows for more variation and complexity in their rRNA genes^8,9^, pathogenic variants in human nuclear rRNA genes are exceedingly rare because of redundancy. Similar to tRNAs, multiple ribosomopathies arise from defects in proteins involved in rDNA transcription, modification, and pre-rRNA processing^10^.

The literature suggests that the majority of human disease-causing tRNA and rRNA sequence variants are found in the mitochondrial DNA (mtDNA), which contains 37 genes, of which 13 are protein-coding, 22 encode for tRNAs and 2 for rRNAs^11,12^. In contrast to the nuclear genome, the mitochondrial genome contains a single copy of each tRNA gene, except for tRNA-Leu and tRNA-Ser, and a single copy of each ribosomal subunit. This lack of redundancy renders mitochondrial RNA genes vulnerable to the effects of sequence mutations. Furthermore, the repair mechanism in mitochondrial DNA is less effective than that in nuclear DNA, increasing its susceptibility to variations and elevating mutation rates. Moreover, the mtDNA is a multicopy genome, therefore one of the key aspects in the biological interpretation of these variants reside in their levels of heteroplasmy, the allelic dosage of each variant^11,12^. Maternal inheritance patterns and tissue-specific expression of mitochondrial genes also add complexity to our understanding of these mutations. Finally, mitochondrial diseases are characterized by significant phenotype and genetic heterogeneity, resulting in a limited availability of well-established genotype-phenotype correlations^13^. The combination of these factors intensifies the likelihood of harmful variants occurring and persisting within mtDNA, not yet identified.

Focusing on mitochondrial tRNAs, 44 single nucleotide variants (SNVs) are currently considered pathogenic in MITOMAP^14^ (i.e., confirmed pathogenic and confirmed likely pathogenic), whereas additional 230 SNVs are likely to be due to an incomplete amount of confirmed evidence from the literature (i.e., reported variants). The associated clinical conditions are developmental, neurological, and metabolic diseases, among others^13^. On the other hand, a recent comprehensive analysis of the human 16S mitochondrial rRNA gene (*MT-RNR2*) identified 145 potentially disruptive mutations. Among these, 64 variants were unique to the source and reported to be potentially pathogenic^15^. To date, MITOMAP contains 65 potentially pathogenic rRNA variants (listed as “reported”), mostly associated with deafness^14^, with the m.1555A>G and m.1494C>T variants of the 12S rRNA (*MT-RNR1)* consistently associated with antibiotic-induced hearing loss^16,17^. Besides their established role in hereditary disorders, the involvement of sequence variants in tRNAs and, more recently, in rRNAs was also proved in complex diseases, such as in cancer, further highlighting the necessity to deeply investigate these noncoding variants^18^. Noncoding RNA dysregulation has been increasingly implicated in cancer progression and therapy response, as shown for example by the association of miR-210-3p expression with poor outcome in docetaxel-treated breast cancer patients^62^.

Considering that noncoding genes span 24% of the total mtDNA length, that 12,014 potential SNVs remain in addition to the 46 known tRNA/rRNA variants, and that variant effect predictors (VEPs) for these variants are either discontinued or lacking, we introduce nAPOGEE (noncoding APOGEE), a VEP specifically developed for mitochondrial noncoding gene variants. It is delivered through MitImpact^19,20^, a collection of genomic, clinical, and functional annotations of all nucleotide changes in the human mitochondrial genes. As APOGEE^21,22^, which was designed for missense variants, nAPOGEE provides scores and probabilities of pathogenicity for five classes of variants, including those of uncertain significance. This makes it compliant with the American College of Medical Genetics and the Association of Molecular Pathology (ACMG/AMP) standards and guidelines^23^.

## Methods

### Training and test datasets

The tRNA and rRNA training datasets were built from “MITOMAP^14^ Disease” and “MITOMAP Polymorphisms” reference files (retrieved in January 2025). From the 1,050 variants in the MITOMAP Disease reference file, indels and variants not localized in tRNA or rRNA genes were removed together with those found in the MITOMAP Polymorphisms file, thereby maintaining only the variants not observed in GenBank, that is, those exhibiting an allele count of zero. The resulting tRNA SNVs were filtered based on the MITOMAP labels “confirmed” or “reported.” The final pathogenic training set included 132 tRNA SNVs (**Dataset 1**). Applying the same filtering strategy to rRNA SNVs, we obtained 22 SNVs, all classified as “reported” in MITOMAP (**Dataset 1**). From the 19,279 variants belonging to the MITOMAP Polymorphisms dataset, we excluded all indels and variants that did not hit tRNA and rRNA genes. SNVs overlapping with the pathogenic dataset were further removed, and the remaining SNVs were further filtered based on allele frequency (AF), maintaining only SNVs with AF ≥ 0.002% in GenBank, as this threshold is indicated as supporting evidence for pathogenicity according to ACMG/AMP guidelines^23^. We obtained a final tRNA benign training set composed of 552 variants and an rRNA training set composed of 1,005 SNVs (**Dataset 2**). To construct reliable test sets for pathogenic and benign tRNA and rRNA, we exclusively considered SNVs that were validated by ClinGen panel experts in accordance with the ACMG/AMP guidelines (repositories were accessed in January 2025). These SNVs were classified as “pathogenic,” “likely pathogenic,” “benign,” or “likely benign,” and were non-overlapping with the training sets. The final tRNA test set comprised 38 pathogenic SNVs (**Dataset 3**) and 33 benign SNVs (**Dataset 4**). For rRNAs, the final test set included only two SNVs classified by ClinGen experts as “likely pathogenic” and “pathogenic” (**Dataset 3**), along with 49 “benign” and “likely benign” SNVs (**Dataset 4**). **Datasets 1**-**4** are available in **Supplementary Table 1** and described in **Extended Data Fig. 1**.

**Extended Data Fig. 1.**
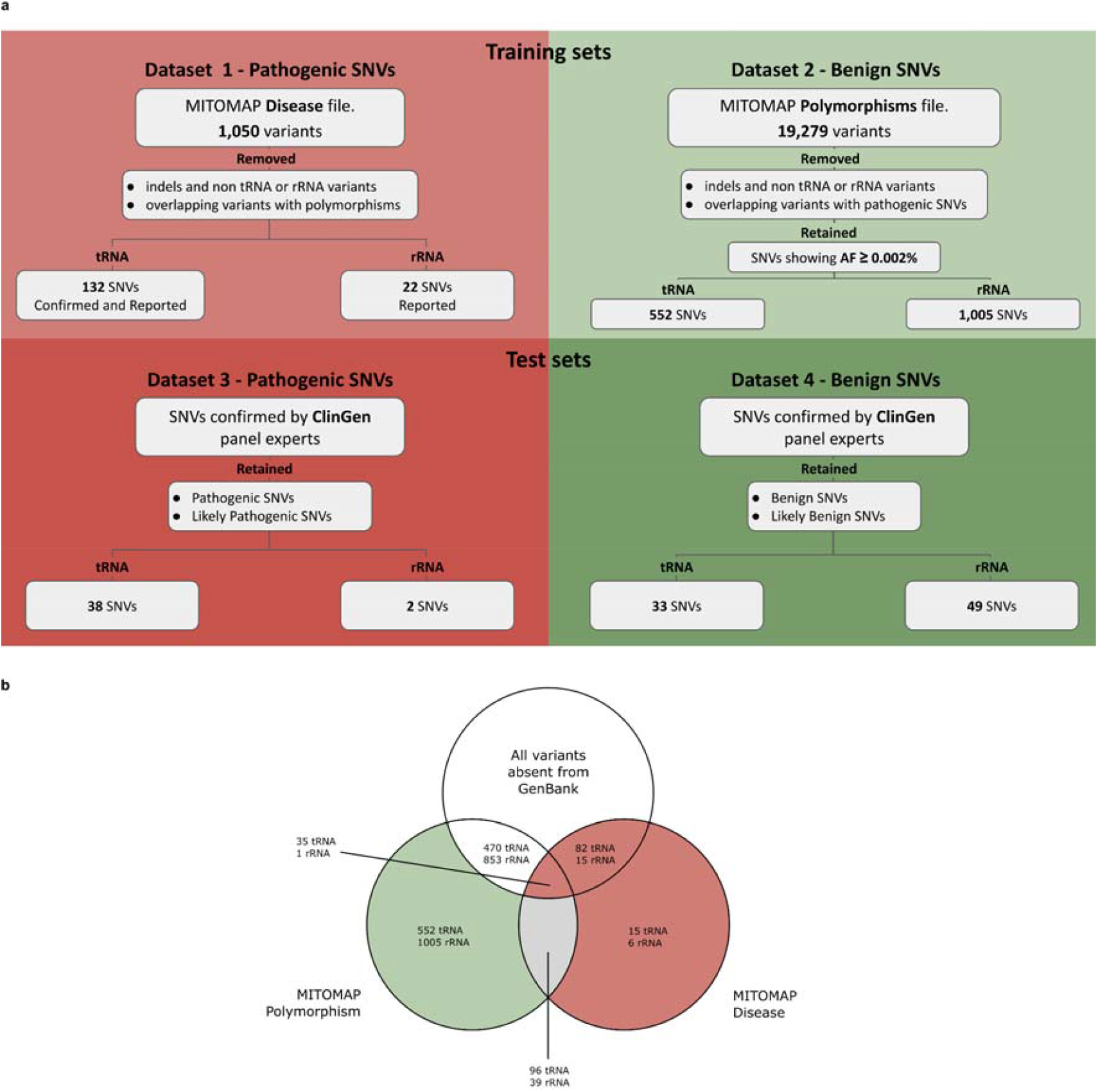
tRNAs and rRNAs datasets. **a**, Schematic representation of the datasets used in this work. Dataset 1 and Dataset 2, used as training sets, were built from MITOMAP Disease and Polymorphisms reference files, respectively. Dataset 3 and 4 included ClinGen curated pathogenic and benign variants classified according to ACMG/AMP guidelines for mitochondrial genome and were used as test sets. The SNVs included in the four datasets are available in **Supplementary Table 1. b**, Inclusion criteria of tRNA and rRNA variants across MITOMAP Disease and Polymorphisms datasets and GenBank; variants included in Dataset 1 are represented in red, while those included in Dataset 2 are shown in green. Variants validated by ClinGen panel experts have not been included in Datasets 1 and 2.

### Features collection

#### Multiple Sequence Alignments

Phylogenetic Multiple Sequence Alignments (MSAs) containing all available Metazoa sequences for all 22 mitochondrial tRNAs were retrieved from tRNAdb^24^ (accessed in March 2023). The taxonomic identifier (taxid) for each species was also recorded. Only 1348 species with complete sets of all 22 tRNAs were retained for further analysis to ensure phylogenetic completeness across taxa. Together with the phylogenetic MSA, we also retrieved the dot-bracket secondary structures associated with each sequence from tRNAdb, integrating structure-function relationship information along with evolution and phylogeny data. We also retrieved the human mitochondrial MSA containing all the 22 aligned human tRNA sequences from tRNAdb.

A phylogenetic MSA for mitochondrial rRNAs was built starting from Metazoa rRNA sequences. We downloaded all Eukariota rRNA sequences as FASTA files for the *MT-RNR1* and *MT-RNR2* genes from MIDORI2^25^ (accessed in February 2025). Among all MIDORI2 sequences, we specifically extracted only those from Metazoan genomes in the NCBI Organelle database, together with their taxonomic lineage. We further filtered out the sequences with at least one missing nucleotide. Only genomes containing one, and only one, copy of both *MT-RNR1* and *MT-RNR2* were retained, resulting in 15,443 genomes across 15,209 distinct metazoan taxon leaves (i.e., species and subspecies). For taxon leaves with multiple genomes, we selected rRNA sequences with the highest similarity to the human reference sequence. The resulting 15,209 *MT-RNR1* and *MT-RNR2* sequences (one per leaf) were aligned using Clustal Omega^26^ to generate distinct MSAs for the two genes.

#### Positional

Each tRNA SNV was mapped onto the human mitochondrial MSA to ensure consistent global nucleotide alignment.

In contrast, mitochondrial rRNAs do not share a human conserved secondary or tertiary structure analogous to that of tRNAs. Therefore, to localize SNVs within rRNA molecules, we utilized spatial coordinates derived from three-dimensional structural complexes. These complexes were generated by submitting wild-type (WT) *MT-RNR1* and *MT-RNR2* sequences to the AlphaFold server^27^. Among the predicted structures, we selected the complex exhibiting the highest structural concordance with known experimental data, as evidenced by its similarity to the deposited PDB entry (6NU2, 1.161 Å RMSD between pruned atom pairs). We opted to use AlphaFold-predicted structures rather than existing PDB models because no available mitochondrial rRNA complex in the PDB contains unmodified WT sequences of both *MT-RNR1* and *MT-RNR2*.

#### Nucleotide Frequencies and Coverage

For each SNV located within the mitochondrial tRNA and rRNA genes, the frequencies of both the reference and alternative nucleotides were computed across the corresponding positions in the phylogenetic MSA. In addition, coverage was quantified as the proportion of sequences exhibiting a non-gap character (i.e., excluding “-”) at the aligned position within the phylogenetic MSA.

#### Unpaired Nucleotide Frequency

We computed the frequency of each tRNA reference unpaired nucleotide using phylogenetically conserved dot-bracket secondary structures retrieved from tRNAdb. This frequency reflects how often a nucleotide localized in a given position remains unpaired across Metazoa species, providing insight into the structural variability of that position across evolutionary lineages.

Due to the lack of public phylogenetic secondary structure data for mitochondrial rRNAs, this analysis could not be extended to *MT-RNR1* and *MT-RNR2*.

#### Phylogenetic Conservation

To evaluate the phylogenetic distances between *Homo sapiens* and other species represented in the MSAs, we used taxonomic lineage information within the Metazoa clade. Lineage data were retrieved from the NCBI Taxonomy database, and phylogenetic relatedness was assessed by comparing the taxonomic hierarchy of each species with that of *Homo sapiens* (TaxID: 9606). The number of hierarchical taxonomic levels shared with *Homo sapiens* was computed for each species, and phylogenetic distance was defined as the reciprocal of this count. Accordingly, smaller values denote a greater evolutionary proximity to humans. Using this metric, we quantified the phylogenetic conservation of each reference nucleotide within tRNA and rRNA genes. Specifically, for each nucleotide position, we identified the phylogenetic distance at which the reference nucleotide was first substituted, either by an alternative nucleotide or a gap, in the MSAs.

#### MSA-based language embedding

To capture the evolutionary context at the nucleotide level, we employed the RNA-MSM language model^28^, pre-trained on phylogenetic RNA MSAs. Embeddings were generated for each SNV in the tRNA and rRNA genes as follows: for each gene, the human reference sequence was first altered to incorporate the SNV of interest; the corresponding MSA was then constructed with the human sequence positioned at the top and the remaining sequences ordered by decreasing similarity; the complete MSA was then embedded using RNA-MSM, yielding a 768-dimensional embedding vector for every position in each sequence (resulting in a tensor of dimensions S × L × 768, where S is the number of sequences, and L is the alignment length, including gap characters). The embedding corresponding to the SNV position in the human sequence was extracted and designated as the final representation of the SNV.

To optimize computational efficiency, each MSA was downsampled to a maximum of 512 sequences using a *diversity-maximizing greedy* algorithm (see *greedy_select* in the RNA-MSM repository), which prioritizes phylogenetic diversity while retaining information content. Given that RNA-MSM was trained on alignments of up to 1024 positions in length, a sliding window strategy was employed for longer alignments, specifically those corresponding to rRNA genes. In these cases, the window was centered on the variant position, with the window size of 256, chosen to balance computational efficiency and contextual completeness.

#### Position Weight Matrix

To quantify nucleotide conservation at each alignment position within mitochondrial tRNA and rRNA sequences, a Position Weight Matrix (PWMs) was constructed using the following procedure: first, a Position Frequency Matrix (PFM) was generated by counting the frequency of each nucleotide (A, C, G, T) at every column of the MSA. The PFM was then normalized to yield a Position Probability Matrix (PPM), representing the relative frequency of each nucleotide at each alignment position. To enable logarithmic transformation and prevent undefined values, a small pseudocount (ε = 10^−15^) was added to all zero-frequency entries in the PPM, ensuring numerical stability without significantly altering the probability distribution. The PWM was subsequently derived by computing the base-2 logarithm of the ratio between the observed nucleotide probability at each position and the expected background probability, which was set to 0.25 under the assumption of a uniform nucleotide distribution. This transformation emphasizes evolutionarily constrained positions, where certain nucleotides occur at frequencies substantially different from those expected by chance.

#### Sequence Entropy

To further characterize nucleotide variability across the aligned mitochondrial tRNA and rRNA sequences, Shannon entropy was computed for each column of the MSA. At each alignment position, the nucleotide composition was treated as a categorical probability distribution. The entropy value was then calculated using the standard Shannon entropy formula:

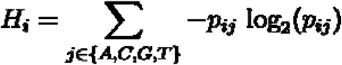

where denotes the relative frequency of base *j* at position *i*. Entropy serves as a quantitative measure of positional uncertainty, with low entropy values indicating high conservation (i.e., reduced variability) and high entropy values reflecting increased sequence heterogeneity.

#### Structural Motif

To assess the structural context of each nucleotide within the mitochondrial tRNA molecules, annotated secondary structure features were incorporated into the analysis. Specifically, positional information for key structural motifs, including bulge loops, stems, hairpin stems, and hairpin loops, was obtained from the MitoVisualize database^29^. Each nucleotide position was then one-hot encoded based on its assignment to these structural elements, thereby enabling integration of secondary structure information into downstream analyses. Due to the absence of a comprehensive and standardized annotation of secondary structure domains for mitochondrial rRNAs, this feature was not extended to rRNA sequences.

#### Secondary structure estimation

The secondary structures of WT tRNAs and all possible variant tRNA sequences resulting from each SNV were predicted using tRNAscan-SE (version 2.0)^30^, with the option *`-M mammal`* designed specifically for mitochondrial tRNA identification. Notably, certain tRNAs lacked nucleotides close to 5’ or 3’ proximities (often the first or last nucleotide) in the predicted secondary structures generated by tRNAscan-SE; therefore, these positions were considered non-paired in subsequent analyses.

For rRNA, secondary structure predictions were performed using the R2DT framework^31^ by applying the templates *d*.*16*.*m*.*H*.*sapiens*.*5* for the 12S rRNA (*MT-RNR1*), and *mHS_LSU_3D* for the 16S rRNA subunit (*MT-RNR2*), as used in the RNAcentral database^32^. These templates were consistently used to generate secondary structures for all potential rRNA SNVs.

#### Change in Gibbs free energy

After structure prediction, the ViennaRNA package (through the command *RNAeval*, version 2.3.3^33^) was employed to compute the change in Gibbs free energy (ΔΔG) associated with each SNV, comparing the thermodynamic stability of the alternative tRNA and rRNA sequences against their corresponding WT references.

#### Secondary structure alteration

To quantify the structural impacts induced by SNVs in tRNAs, we systematically analyzed and compared the secondary structures predicted by tRNAscan-SE for WT versus alternative sequences. Structural changes were quantified using the dot-bracket Hamming distance metric between the predicted structures for each alternative sequence, relative to its WT counterpart. Given the substantially greater lengths of rRNA molecules (954 bp for *MT-RNR1* and 1558 bp for *MT-RNR2*) compared to tRNAs (approximately 70 bp), we did not observe relevant structural changes caused by SNVs in rRNAs. Consequently, structural alteration analysis was exclusively performed for tRNAs and was not extended to rRNAs.

#### Phylogenetic scores

Evolutionary conservation at each genomic position was further quantified using precomputed PhyloP (100 vertebrates)^34^ and PhastCons (100 vertebrates)^35^ conservation indices for both tRNAs and rRNAs.

#### Regional tolerance to genetic variation

To assess regional tolerance to genetic variation within mitochondrial RNA regions, we used the recently described mtDNA local constraint (MLC) scores^36^, which provide a positional measure of constraint throughout the mtDNA. Furthermore, for tRNA SNVs, the observed-to-expected variant frequency ratio (OEUF) scores, derived as conservative metrics of evolutionary constraints from previously published datasets, were employed^36^. These OEUF scores, computed for 75 unique phylogenetically conserved positions in tRNA secondary structures^37^, presented gaps at five positions (m.7474 and m.7492 in *MT-TS1*, m.12252 in *MT-TS2*, and m.8343 and m.8351 in *MT-TK*). Missing values were imputed using the median OEUF score.

#### PTM site distance features

To evaluate the potential impact of SNVs on post-transcriptional modifications (PTMs), we calculated both linear and spatial distances from each SNV to the closest known PTM sites. PTM sites in tRNAs were sourced from the MitoVisualize database^29^, whereas rRNA PTM sites were obtained from the established literature^38^. Linear distances were calculated directly along the nucleotide sequence, whereas spatial distances were computed within the context of the predicted secondary structure modeled as a graph. In this graphical representation, nucleotides constitute nodes and base pairings, excluding pseudoknots, are represented as edges, enabling the computation of the shortest paths between SNVs and PTM sites.

### Training and hyperparameter tuning

#### Preprocessing

A systematic preprocessing pipeline was developed to address the feature selection, scaling, and dimensionality reduction tasks (**Fig. 1**). The pipeline consists of the following sequential steps:

**Fig. 1.**
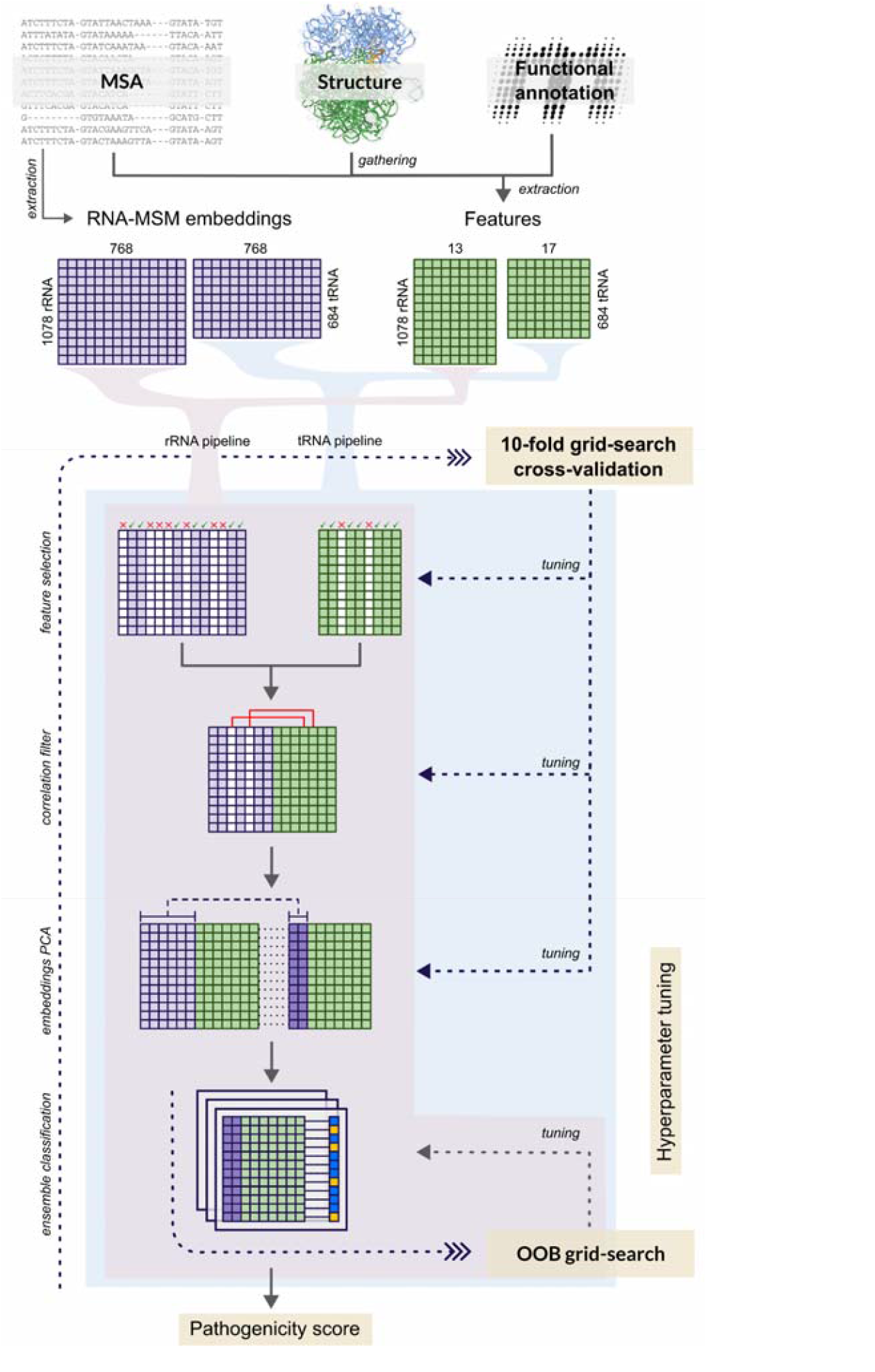
tAPOGEE and rAPOGEE pipelines. The scheme represents the full nAPOGEE pipeline for tRNA and rRNA SNVs classification. The same pipeline was used to obtain 2 different classification models: tAPOGEE for tRNAs and rAPOGEE for rRNAs. The pipeline takes several features derived from MSA, structure and functional annotation in input. Contextually, MSA is also used as input for the RNA-MSM language model to retrieve embedding channels for SNVs. Several preprocessing steps occur before the proper classification: selection of informative features and channels; dropping of embedding channels that show high correlation with non-embedding features; dimensionality and collinearity reduction of the remaining channels through PCA. Finally, an out-of-bag (OOB) grid search is performed to tune a balanced ensemble classifier, lastly trained with the best hyperparameters. Also preprocessing steps require some hyperparameters to be tuned: this has only been done for tAPOGEE via 10-fold grid search cross validation, while for rAPOGEE, the same hyperparameters resulting from tAPOGEE grid search were used.

□ *Supervised tree-based feature selection*: A decision tree classifier employing balanced class weighting, was implemented to select biologically relevant features according to their Gini importance scores. The hyperparameters of the classifier were optimized using a grid-search cross-validation (CV) approach. Decision trees were chosen for their ability to model nonlinear and localized relationships between features and the target class as well as their robustness against feature collinearity.
□ *Supervised FDR-based embedding channel selection*: A False Discovery Rate (FDR)-based method was used to identify RNA-MSM embedding channels (features) associated with SNV classification, as determined by t-test conducted on the training sets (**Datasets 1** and **2** for tRNAs and **Datasets 1-4** for rRNAs). Starting from 768 original embedding channels, only those demonstrating significant associations were retained with the adjusted p-value threshold for significance tuned during the grid-search CV.
□ *Unsupervised correlation-based feature filtering*: A correlation-based filtering strategy was implemented to discard redundant embedding channels. Spearman correlation coefficients were computed between the RNA-MSM embedding channels and previously selected biological features (obtained from the supervised decision tree classifier). Channels exhibiting absolute correlation coefficients (either positive or negative) above a predefined threshold, optimized via grid-search CV, were excluded to reduce redundancy.
□ *Standard Scaling*: All retained features were standardized by transforming them to have zero mean and unit variance.
□ *PCA*: To further reduce collinearity among the remaining RNA-MSM embedding channels, dimensionality reduction was performed using Principal Component Analysis (PCA). The optimal number of principal components was fine-tuned via a grid-search CV.

This preprocessing pipeline was applied to both tRNA (tAPOGEE) and rRNA (rAPOGEE) classification tasks.

#### Preprocessing hyperparameters tuning

We conducted a comprehensive 10-fold grid search CV across the entire tAPOGEE pipeline to refine the preprocessing hyperparameters in order to maximize the average Area Under the Receiver Operating Characteristic curve (AUROC) across the 10 folds. Folds for CV were randomly generated to ensure an equitable distribution of pathogenic variants among each fold. A fixed random seed was employed to guarantee the reproducibility of the results and consistency in the hyperparameter tuning for each ensemble classifier. The specific preprocessing hyperparameters investigated through this grid search, mentioned in the preprocessing section, are summarized in **Supplementary Table 2**. Because of the limited number of rRNA SNVs in the pathogenic test set (**Dataset 3**), we did not perform preprocessing hyperparameter tuning for rAPOGEE. Therefore, we used the same hyperparameters resulting from tAPOGEE tuning.

#### Ensemble Classifiers

We systematically evaluated multiple ensemble-learning methods to serve as core classifiers in our pipeline. The classifier that achieved the highest AUROC during validation was selected. The candidate ensemble classifiers implemented for tAPOGEE were:

□ *Balanced Random Forest*: An ensemble comprising 500 decision trees. To mitigate the class imbalance, each tree was trained on a balanced subset containing 75 pathogenic (from **Dataset 1**) and 75 benign variants (from **Dataset 2**) drawn without replacement.
□ *KNN Balanced Bagging*: An ensemble consisting of 500 K-nearest neighbor (KNN) classifiers. Each base learner was trained on a bootstrap sample equal in size to the original dataset and subsequently subsampled to maintain a balanced (1:1) ratio of pathogenic to benign variants.
□ *SVM Balanced Bagging*: An ensemble of 500 support vector machine (SVM) classifiers with radial basis function (RBF) kernels. The training of each SVM utilized the same balanced bootstrapping approach as that described for the KNN ensemble.

For rAPOGEE, we applied a similar ensemble framework adjusted to accommodate a significantly greater class imbalance and a smaller number of pathogenic variants; for this reason, we used all the **Datasets 1**-**4** for training. Moreover, Balanced Random Forest trees were trained on subsets consisting of 22 pathogenic (from **Dataset 1** and **3**) and 22 benign variants (from **Dataset 2** and **4**), drawn without replacement. To ensure robust predictions and guarantee that each benign variant appeared in multiple base learners, the number of base estimators for rAPOGEE was increased to 4,096 (i.e., 2^12^). This approach aimed to enhance the reliability of out-of-bag (OOB) predictions, particularly for the infrequent pathogenic class.

#### Grid Search with OOB Estimation

Given that all candidate core classifiers are ensemble-based methods, we employed an OOB grid search strategy to optimize the hyperparameters of each classifier. The OOB approach provides a direct estimate of the model performance without requiring a separate validation set, thereby increasing the computational efficiency. Hyperparameter optimization targeted the maximization of the AUROC. In **Supplementary Table 3**, we describe the specific hyperparameters tuned for each ensemble classifier.

The OOB grid search is implemented as the final stage of both tAPOGEE and rAPOGEE pipelines (**Fig. 1**). This design enabled the ensemble classifiers to directly use the features derived from the preceding preprocessing steps as inputs for hyperparameter optimization.

#### Feature importance estimation

Because the SVM Balanced Bagging was found to be the best classifier for the tAPOGEE pipeline and it does not inherently provide directly interpretable estimates of the feature importance, we employed the SHapley Additive exPlanations (SHAP) permutation explainer^39^ with default parameters to estimate the importance of each feature. This estimation was achieved by calculating the expected absolute SHAP value for each feature on the independent tAPOGEE test datasets (**Datasets 3** and **4**) in accordance with the methodology proposed by the original SHAP authors. For each test variant, the SHAP values of the principal components (PCs) derived from the RNA-MSM embeddings were summed to yield an aggregate measure representing the overall contribution of RNA-MSM embeddings to variant classification. Consequently, feature importance assessment reflects the importance of collective RNA-MSM embeddings rather than that of individual PCs.

In contrast, the rAPOGEE classifier uses a Random Forest approach, inherently enabling direct estimation of feature importance quantified as the mean normalized reduction in entropy (information gain) obtained across all trees in the ensemble.

### Comparative Analysis with Existing mt-tRNA VEPs

Given the current absence of VEPs specifically tailored to rRNAs, we benchmarked tAPOGEE performance and compared it with that of the existing mitochondrial tRNA VEPs. Specifically, we compared tAPOGEE to PON-mt-tRNA^40^ and Mitotip^41^.

#### PON-mt-tRNA

We downloaded PON-mt-tRNA precomputed predictions for all 4521 possible human tRNA SNVs (and, specifically, PON-mt-tRNA ML probability). To directly compare the PON-mt-tRNA and tAPOGEE performances while preventing any training bias, we used a subset of **Datasets 3** and **4** as a test set, specifically dropping variants overlapping with the PON-mt-tRNA training set^42^. We repeated the same test after retraining the tAPOGEE model on the PON-mt-tRNA training set^42^.

#### MitoTIP

MitoTIP^41^ is currently indicated as one of the computational evidence to assign pathogenic/benign criteria to tRNA variants and is recommended by the ACMG/AMP standard guidelines^23^. We downloaded MitoTIP precomputed predictions for all possible tRNA SNVs, along with its training set variants. As already done for PON-mt-tRNA, we first compared tAPOGEE and MitoTIP performances on **Datasets 3** and **4** purged from MitoTIP training set variants. Subsequently, we retrained tAPOGEE on the MitoTIP training set and tested both tools on **Datasets 1** and **2**, after removing the MitoTIP training set variants. As last experiment we performed a 5-times repeated 10-fold stratified CV of tAPOGEE on **Dataset 1** and **2**: for each CV iteration we evaluated tAPOGEE and MitoTIP on variants excluded from both training folds and MitoTIP training set, ensuring that no variant used during evaluation had been seen by either tAPOGEE or MitoTIP during training.

### Predictions and classification of tRNA and rRNA variants according to ACMG/AMP guidelines

We adopted the Bayesian inference approach as previously done by Bianco et al.^22^ to compute the posterior probability of pathogenicity of tRNA and rRNA SNVs. Specifically, we estimated the likelihood distributions of nAPOGEE scores separately to reliably classify benign and pathogenic SNVs using the variants contained within **Dataset 4** (benign and likely benign revised by ClinGen experts) and **Datasets 1** and **3** (MITOMAP- and ClinGen-selected pathogenic variants), respectively. Benign variants from the training dataset (**Dataset 2**) were intentionally excluded from this analysis to prevent the inclusion of false positives resulting from the allele frequency-based strategy used to build this dataset. Given that the nAPOGEE scores are bounded between 0 and 1, their distributions are modeled using zero-one inflated beta distributions. Furthermore, we used OOB predictions for variants originating from the training set (**Dataset 1**) to mitigate potential bias in the posterior probability estimations. We adopted a prior probability of pathogenicity of 0.10, deriving the marginal distribution of nAPOGEE scores by weighting and summing both likelihood distributions according to their respective prior probabilities (0.90 benign, 0.10 pathogenic).

The Bayesian procedure described above was conducted independently for tRNA and rRNA SNVs, resulting in distinct estimates tailored to each RNA type.

### Spatial analysis of pathogenicity

To evaluate spatial autocorrelation in tRNA variant pathogenicity scores, we quantified global spatial autocorrelation using Moran’s index and identified regional contributions through Local Indicators of Spatial Association (LISA). Pairwise nucleotide distances within each mitochondrial tRNA were computed based on the shortest-path distances within their secondary structures, following the methodology previously used to identify the nearest PTM sites. To avoid spatially induced bias and isolate feature-driven pathogenicity predictions, positional-independent predictions were generated by excluding ensemble classifiers trained on positional features (i.e., the human MSA position and nearest PTMs).

For each tRNA consisting of *L* nucleotides and *3L* variants, the LISA statistic for variant □ was computed as follows:

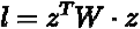

where □ is a standardized vector of tAPOGEE scores for all SNVs mapping onto the specific tRNA under investigation, and □ is a normalized spatial-weight matrix reflecting nucleotide proximity within the secondary structure, defined as:

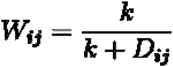

where *D*_*ij*_ represents the shortest distance between the variant *i* and *j* and *k* = 1. The resulting *l*_*j*_ ∈ ℝ ^3*L*^ is a vector and each *l* ∈ *l* thus quantifies the coherence between the pathogenicity score of variant □ and scores of variants in its immediate spatial vicinity.

The global spatial autocorrelation within each tRNA was summarized using Moran’s index (I):

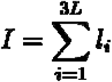

where □ ranges from -1 to 1, with positive values indicating spatial clustering of similar pathogenicity scores and negative values indicating spatial dispersion.

To statistically assess whether the observed Moran’s index or LISA values significantly deviated from the null hypothesis of no autocorrelation, we performed 10,000 permutations of *z*, correcting p-values using the Benjamini-Hochberg method (denoted as *q-values*). Since each nucleotide is associated with three overlapping SNVs, we assigned a single LISA value per nucleotide by averaging the corresponding three SNV-level LISA values and conservatively retained the highest corrected p-value among them.

We repeated the same procedure for the rRNA complex (including both rRNA subunits) after filtering out all rAPOGEE estimators using at least one positional feature (i.e., 3D coordinates x, y, z, and the closest PTMs) and defining *W* in function of the Euclidean distance of SNVs on the rRNA complex. Moreover, since we were working on physical distances, we set *k* = 10.85Å as the ideal C1′–C1′ distance for Watson– Crick base pairs.

### Molecular Dynamics Simulation

Three-dimensional atomic models of both WT mt-rRNAs (see positional features description), together with their respective mutant structures, were obtained using the AlphaFold server^27^, and the 12S and 16S complexes were recreated by superimposition using ChimeraX^43^. In contrast, we employed the RNA Composer web server^44^ to generate atomic models of WT and mutant mt-tRNAs, submitting their primary sequences and corresponding secondary structure annotations retrieved from tRNAdb^24^. Each model was then embedded in a TIP3P-solvated water box using a CHARMM-GUI web interface.

Classical MD simulations were performed for 200 ns in triplicate using Amber20 with the Amber ff14SB force field^45,60,61^. The resulting trajectories were analyzed using specific geometric criteria. Root Mean Square Deviation (RMSD) and Root Mean Square Fluctuation (RMSF) profiles were computed using the MDAnalysis Python library. Secondary structures in dot-bracket notation were derived for each trajectory frame using the X3DNA-DSSR tool^46^, and consensus structures were computed. Finally, dynamic secondary structure annotations were generated using the SEC_STRUCTURE function of the Barnaba toolkit^47^.

### Permutation-Based Significance Testing

For all reported AUROC and Average Precision (AP) values, statistical significance was assessed against a null distribution obtained via 10,000 random label permutations. A one-sided empirical p-value was computed as the proportion of permutations yielding a score greater than or equal to the observed one. All reported performances correspond to cases where p-value < 0.05.

## RESULTS

### Overview of Model Development

We developed two specialized machine learning frameworks for the classification of pathogenicity of noncoding mitochondrial variants, tAPOGEE for mt-tRNA variants and rAPOGEE for mt-rRNA variants (**Fig. 1**). For general reference, we use the term nAPOGEE (“noncoding APOGEE”) to denote the unified framework when broadly referring to the classification of noncoding mitochondrial variants. Both pipelines were rigorously trained and validated on carefully curated variant datasets and extensively annotated with biologically informative features, as described in detail in the Methods section.

### Construction of Variant Datasets

For the tAPOGEE pipeline, we assembled a training dataset comprising 132 pathogenic and 552 benign tRNA SNVs (**Dataset 1** and **2, Extended Data Fig. 1, Supplementary Table 1**). Benign variants were sourced from the “Polymorphisms” repository of MITOMAP^14^, whereas pathogenic variants were identified by combining MITOMAP’s “reported” and “confirmed” disease-associated variant lists. To evaluate the generalizability of the tAPOGEE model rigorously, an independent test dataset was constructed using variants annotated by ClinGen, comprising 38 pathogenic or likely pathogenic variants and 33 benign or likely benign variants (**Dataset 3** and **4**).

For rAPOGEE, the training dataset included 22 reported pathogenic and 1,005 benign rRNA SNVs obtained from MITOMAP (**Dataset 1** and **2**). Due to the limited availability of known pathogenic rRNA variants, the independent ClinGen dataset was inadequate for external validation, containing only two pathogenic or likely pathogenic variants and 49 benign or likely benign variants (**Dataset 3** and **4**). Consequently, we integrated these ClinGen-derived variants directly into the training set, thereby optimizing the available information for model training. The class imbalance observed in rRNA data reflects the paucity of dedicated tools for predicting variants located in these genes^23^, which are generally discarded, resulting in their relative rarity compared with benign variants, thus posing inherent challenges for their accurate classification.

### Model Training, Selection, and Evaluation

For the tAPOGEE pipeline, preprocessing hyperparameters were rigorously optimized through a comprehensive grid-search cross-validation (CV), leveraging a relatively large and balanced variant training dataset. Conversely, for the rAPOGEE pipeline, no preprocessing hyperparameter tuning was performed due to the limited number of rRNA SNVs in the pathogenic test set. Therefore, the same hyperparameters optimized for tAPOGEE were reused.

In the classification stage, three distinct ensemble methods were systematically evaluated within each pipeline: *Balanced Random Forest, Balanced K-Nearest Neighbor (KNN) Bagging*, and *Balanced Support Vector Machine (SVM) Bagging*. Hyperparameter optimization for each classifier was performed using the out-of-bag (OOB) predictions during the grid-search. The final model selection was based on the highest OOB AUROC value. Specifically, tAPOGEE selected the Balanced SVM Bagging model, achieving a grid-search CV AUROC of 0.83 and an OOB AUROC of 0.84 (**Fig. 2a**). rAPOGEE selected the Balanced Random Forest model, which achieved a grid search OOB AUROC of 0.98 (**Fig. 2b**). To assess the generalization performance, the final tAPOGEE classifier was evaluated on an independent ClinGen-curated test set, achieving an AUROC of 0.93 and an Average Precision (AP) of 0.95 significantly higher than the 0.54 expected AP. Due to the absence of an adequately sized independent test dataset for rRNA variants, rAPOGEE performance was assessed via a 24-fold stratified CV strategy, systematically excluding one pathogenic variant per fold. Using this rigorous approach, rAPOGEE achieved an AUROC of 0.87 (95% CI: 0.78 - 0.95, **Fig. 2b**) and an AP of 0.49 vs the expected 0.02 random AP (95% CI: 0.04 - 1.00).

**Fig. 2.**
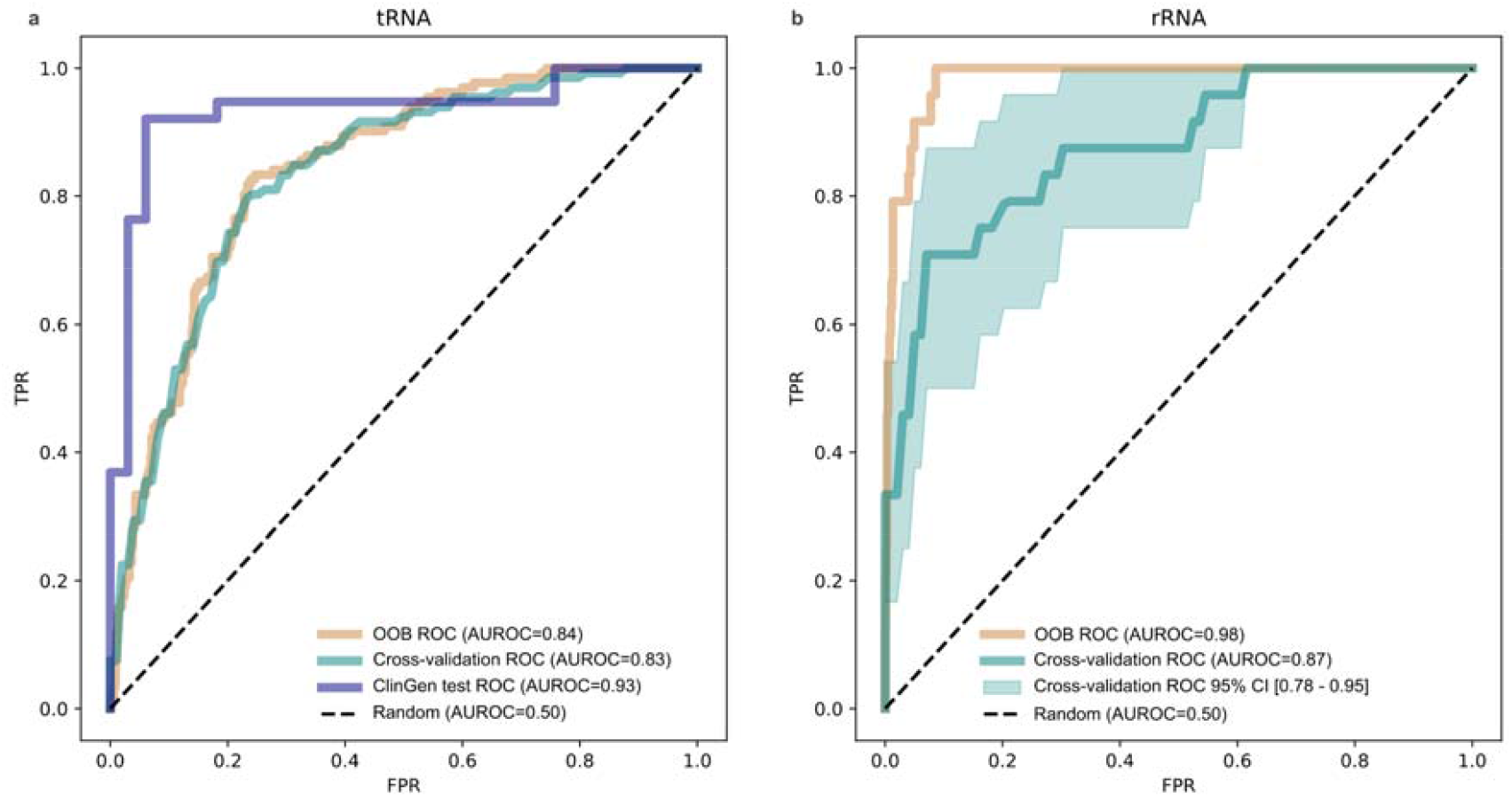
ROC performances of the tAPOGEE and rAPOGEE models. **a**, ROC curves for tAPOGEE evaluated using: out-of-bag (OOB) estimation during classifier tuning (light orange, AUROC□=□0.84), 10-fold cross-validation (CV) applied for preprocessing hyperparameters tuning (teal, AUROC□=□0.83), and final independent evaluation on the ClinGen test set (**Datasets 3** and **4**, dark blue, AUROC□=□0.93). **b**, ROC curves for rAPOGEE using OOB estimation during ensemble hyperparameter tuning (light orange, AUROC□=□0.98) and 24-fold CV used for model evaluation due to the lack of an external test set (teal, AUROC□=□0.87). The shaded area represents the 95% confidence interval of the CV ROC (0.78□-0.95). The dashed diagonal line represents the performance of a random classifier (AUROC□=□0.50).

### Feature Contribution Analysis

The initial set of features considered for developing tAPOGEE and rAPOGEE underwent a feature selection process during the preprocessing, using a decision tree classifier for features derived from phylogenesis, structure and functional annotation (threshold set at 0.01 importance, as indicated in **Supplementary Table 2**), and a specifically designed filtering strategy for RNA-MSM embedding channels (**Fig. 1**, Gini importance threshold set at 0.01, **Supplementary Table 2**). As a result, only the most relevant features were retained for model training, while others were discarded.

To gain insight into the predictive mechanisms employed by our classifiers, we conducted detailed feature importance analyses tailored to each model’s structure. As detailed in the Methods section, for the tAPOGEE model, feature contributions were evaluated using SHapley Additive exPlanations (SHAP) (**Extended Data Fig. 2a**). Conversely, for the rAPOGEE model, we directly retrieved the feature importance from the Random Forest model (**Extended Data Fig. 2b**).

Notably, in tRNA classification the features capturing sequence evolutionary conservation were key contributors. PhastCons accounted for 13% of total importance and the Phylogenetic Conservation feature explained 11% of overall importance. Differently, in rRNA classification, the structural features, i.e., the 3D coordinates (x, y, and z coordinates) and closeness to a post-transcriptional modification (PTM) site played a prominent role (19% and 9% of total importance). For both classification pipelines, MLC constraint score represented a very useful feature (14% and 8.5% of total importance for tRNAs and rRNAs, respectively), indicating that these noncoding molecules contain regions under a strong purifying selection, which are difficult to capture through other features.

Strikingly, for both tRNA and rRNA variants, RNA-MSM embedding channels emerged as highly informative features, accounting for 21□% and 31□% of total importance, respectively. Specifically, among the 768 available channels, 158 and 28 were identified as informative for tAPOGEE and rAPOGEE, respectively, but only 82 and 21 were retained as they did not show collinearity with the other features (see Methods). These results suggest both the substantial contribution of RNA-MSM for SNVs classification and its ability to capture latent properties of mitochondrial noncoding RNAs that are not fully represented by biological and structural features conventionally used to describe variants located in these RNAs.

**Extended Data Fig. 2.**
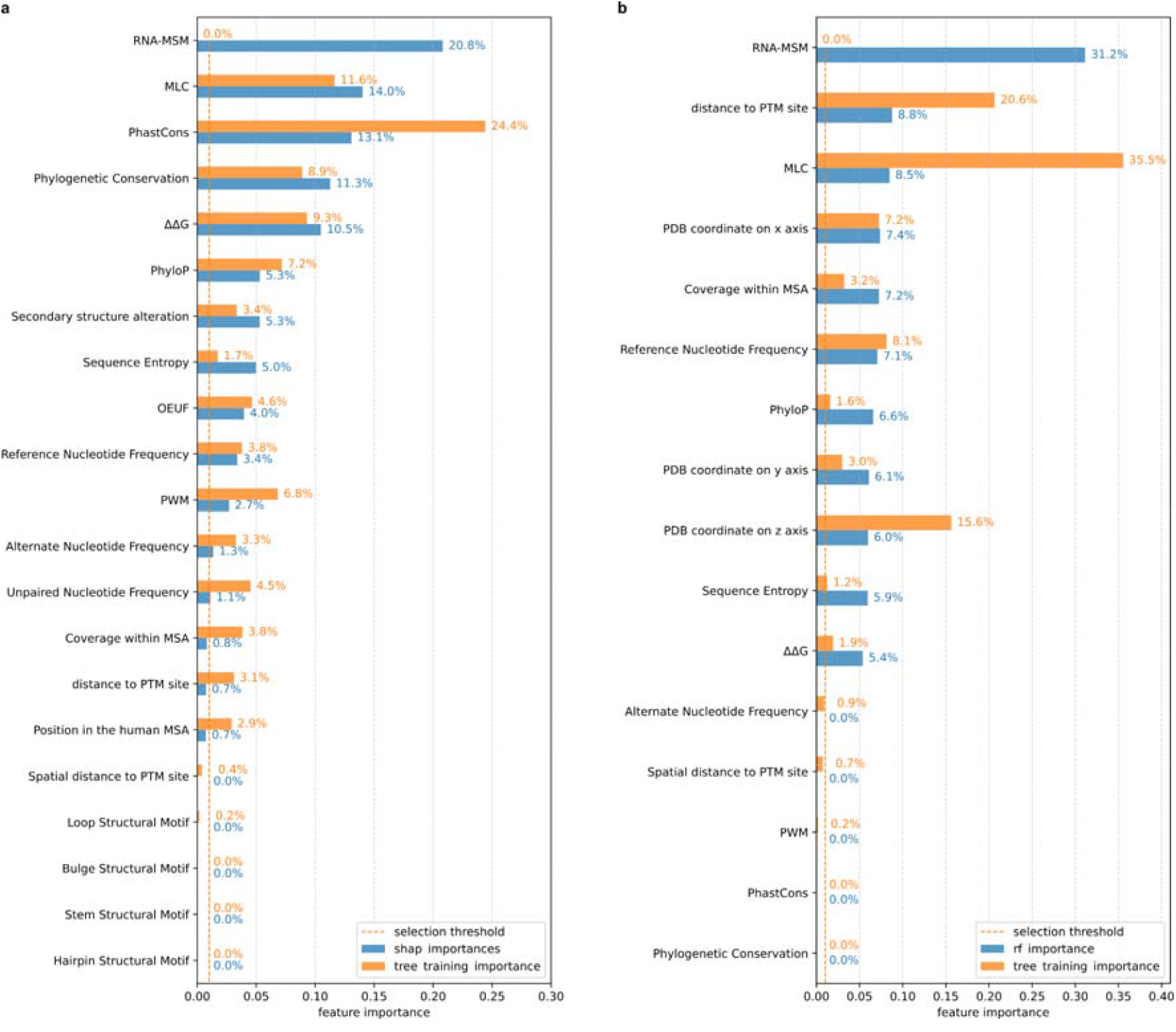
tAPOGEE and rAPOGEE feature importance. As detailed in the Methods section and represented in **Fig. 1**, the initial set of features used to develop tAPOGEE and rAPOGEE went through a feature selection step, performed using a specifically designed filtering strategy for RNA-MSM and a decision tree classifier for all the other features included in this work. While RNA-MSM feature importances could not be represented (due to the different feature selection procedure), the biological and structural decision tree feature importances are reported in **a** and **b** in orange (for tRNAs and rRNAs, respectively). For the features selected in the tAPOGEE and rAPOGEE models we evaluated their importances using SHAP for tRNAs (in blue, **a**) and retrieving their values directly from Random Forest model for rRNAs (in blue, **b**); the feature importance belonging to RNA-MSM embeddings used in the model training was aggregated to generate the figure.

### Comparative Evaluation with PON-mt-tRNA and MITOTIP

We evaluated the predictive performance of tAPOGEE against the existing mitochondrial tRNA variant effect predictor PON-mt-tRNA^40^ using a subset of the ClinGen-curated test dataset after excluding all variants previously employed in PON-mt-tRNA’s training set. This resulted in a carefully curated comparative test set containing 31 variants (9 pathogenic and 22 benign). In this test set, tAPOGEE achieved an AUROC of 0.92 and an AP of 0.93, surpassing the expected random AP baseline of 0.29. In comparison, PON-mt-tRNA attained an AUROC of 0.80 and an AP of 0.76.

To further assess the robustness and generalizability of tAPOGEE, we retrained the model using exactly the same variant dataset originally used by PON-mt-tRNA (containing 146 variants) and tested on the same 31 variants, as above described, deliberately omitting additional hyperparameter tuning to minimize computational complexity. Remarkably, even under these conditions, tAPOGEE achieved an AUROC of 0.90 and an AP of 0.81, again exceeding PON-mt-tRNA’s performance (see **Table 1**).

**Table 1.** Comparison between tAPOGEE and PON-mt-tRNA performances.

|  | **AUROC** | **AP** |
| --- | --- | --- |
| **PON-mt-tRNA** | 0.80 | 0.76 |
| **tAPOGEE** | 0.92 | 0.93 |
| **tAPOGEE trained on PON-mt-tRNA training set** | 0.90 | 0.81 |
| **Random Classifier** | 0.50 | 0.29 |

We repeated the same procedure to compare tAPOGEE with MitoTIP^41^ obtaining a subset of 19 SVNs from **Dataset 3** and 13 from **Dataset 4** for a total of 32 test variants. As expected, since ClinGen classified variants are curated and have much support in the literature, MitoTIP exhibits a 0.99 AUROC and 0.99 AP overperforming tAPOGEE, which shows a 0.90 AUROC and a 0.93 AP (see **Table 2**). Therefore, we retrained tAPOGEE on the MitoTIP training set. Testing both VEPs on a less characterized dataset, consisting of **Datasets 1** and **2**, purged of MitoTIP training set, tAPOGEE showed better results: MitoTIP achieved an AUROC of 0.75, AP of 0.64, tAPOGEE achieved an AUROC of 0.77 and AP of 0.73. This indicates that tAPOGEE prioritizes novel and uncharacterized variants (including those of uncertain significance, VUS) more effectively, whereas MitoTIP’s predictive performance depends heavily on prior literature annotations and population frequency data.

**Table 2.** Comparison between tAPOGEE and MitoTIP performance. Both **Datasets 3** and **4** and **Datasets 1** and **2** were purged of MitoTIP training set variants, as indicated in the column “Considered variants”, which were used to test the different pipelines listed on the right.

| **Datasets** | **Considered variants** | **Tools and criteria used to test** | **AUROC** | **AP** |
| --- | --- | --- | --- | --- |
| **3 and 4** | 32<br>(19 P + 13 N) | **MitoTIP** | 0.99 | 0.99 |
|  |  | *tAPOGEE* | 0.90 | 0.93 |
|  |  | *Random Classifier* | 0.50 | 0.60 |
| **1 and 2** | 230<br>(112 P + 118 N) | *MitoTIP* | 0.75 | 0.64 |
|  |  | *tAPOGEE trained on MitoTIP training set* | 0.77 | 0.73 |
|  |  | *cross-validation of tAPOGEE on Dataset 1 and 2* | 0.78 | 0.77 |
|  |  | *Random Classifier* | 0.50 | 0.49 |

To further test tAPOGEE, we performed an additional cross-validation run on **Datasets 1** and **2**, which included only variants not overlapping with the MitoTIP training set and the current tAPOGEE training fold. Under this evaluation setting, tAPOGEE achieved even higher performance, with an average AUROC of 0.78 and AP of 0.77, significantly outperforming MitoTIP according to both AUROC (Wilcoxon signed-rank test, two-sided p = 0.025) and AP (p < 0.0001).

### Structural mapping of nAPOGEE scores

We then considered the localization of nAPOGEE scores along tRNAs and rRNAs separately (**Fig. 3**). Considering tRNAs, we found that *MT-TM* showed the highest nAPOGEE median score, probably reflecting its biological role as the gene encoding the tRNA-Met, which is crucial for translation initiation (**Fig. 3c**). When we considered the average nAPOGEE scores on the phylogenetically conserved tRNA secondary structure^37^, we found that, as expected, anticodon stem and loop regions tend to accumulate variants with higher scores (**Fig. 3a**). We also observed that T and D stems are more likely to accumulate deleterious variants differently from their respective loops, which are more likely to be prone to neutral variants (**Fig. 3a**). Interestingly, as already reported by Lake et al.^36^, we found that D-stem accumulated variants with the highest nAPOGEE scores (average nAPOGEE =0.65), while T-loop accumulated variants with lower scores (average nAPOGEE =0.27), reflecting the different constraints in these regions. Consequently, looking at single nucleotides, we observed that those accumulating the highest nAPOGEE scores were localized in the D-stem region (above all, in the positions 10 and 24, **Fig. 3a**), which has a fundamental role in forming the tertiary tRNA structure^36^.

**Fig. 3.**
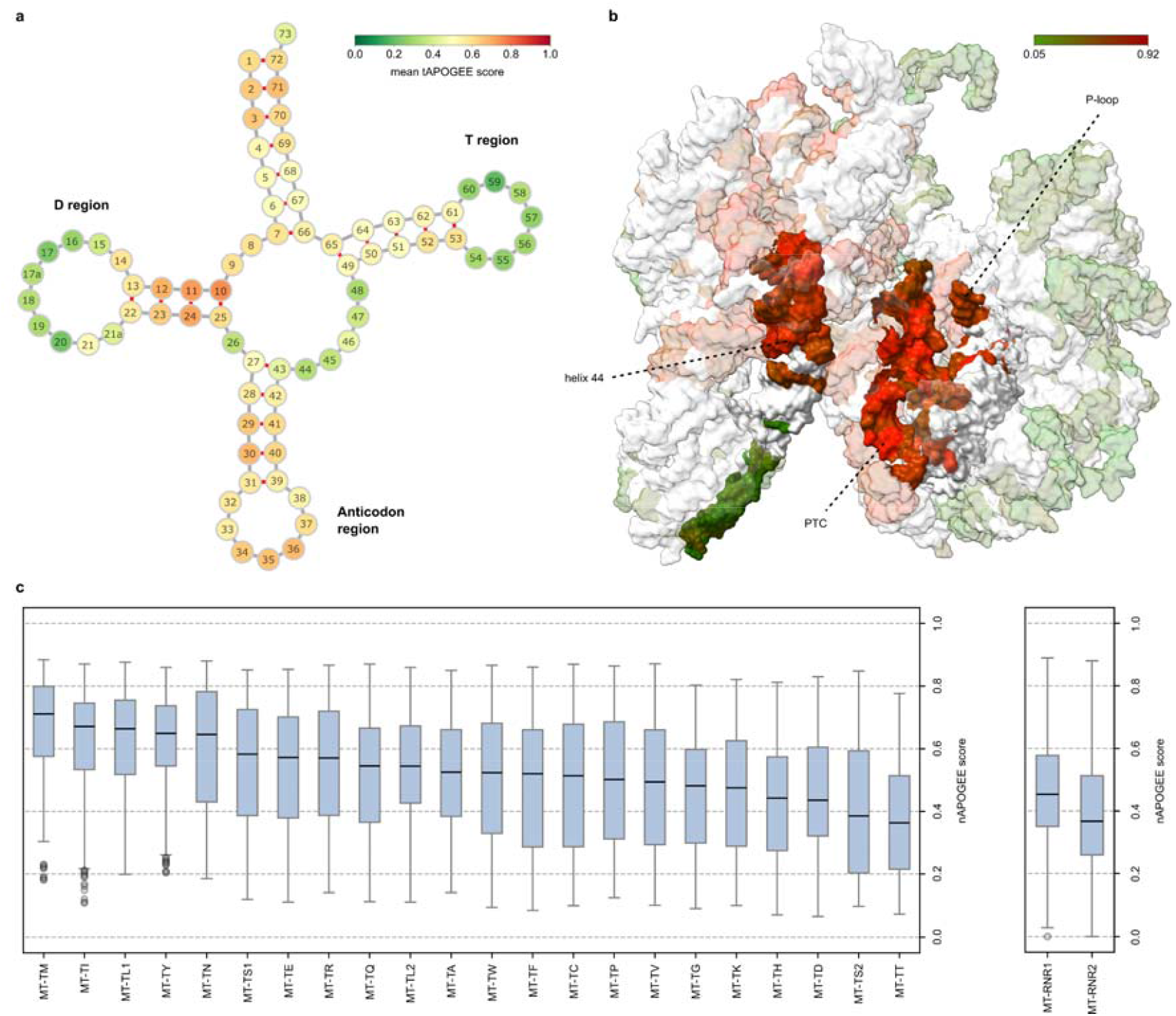
Residue-dependent nAPOGEE score. **a**, nAPOGEE scores have been mapped on the phylogenetically conserved tRNA secondary structure, consisting of 75 unique positions, taken from^37^. Scores have been mediated for each position of the structure and represented in a color scale from green to red, i.e. from lower to higher scores. **Extended Data Fig. 3** shows scores for individual tRNAs. **b**, nAPOGEE scores mapped on the rRNA complex, obtained through AlphaFold; only residues showing significantly positive LISA values have been colored according to their average nAPOGEE scores, from green to red. In the figure we evidenced functional regions, in which significant scores were found, i.e., P-loop, peptidyl transferase center (PTC) region and H44 helix. A 3D map of significantly positive LISA nAPOGEE scores for the rRNA complex can be found at [https://github.com/mazzalab/nAPOGEE/blob/main/rAPOGEE/checkpoints/lisa.html]. **c**, nAPOGEE scores across noncoding genes. Boxplots were generated to represent the interquartile range (IQR), defined as the interval between the 25^th^ percentile (Q1) and the 75^th^ percentile (Q3). The central line within each box denotes the median (Q2), while the whiskers extend to Q1 – 1.5×IQR (lower bound) and Q3 + 1.5×IQR (upper bound).

We also investigated the spatial autocorrelation of nAPOGEE scores for each tRNA (**Extended Data Fig. 3**). Hence, we excluded estimators based on variant localization and, therefore, did not consider information on the alignment and their distance to the PTM sites. We observed significant positive spatial correlation for nAPOGEE score for all tRNAs secondary structures (permutation *q-value*<0.0001 for every tRNA) with an average Moran’s index of 0.17 (ranging from 0.11 to 0.23); complete data can be found in **Supplementary Table 4**. Local indicators of spatial association (LISA) of positional bias-free nAPOGEE scores obtained from this analysis pointed out that almost all tRNAs had a significant spatial autocorrelation of nAPOGEE low scores in T-loop regions (LISA > 0 and *q-value* < 0.05, **Extended Data Fig. 3**). Taken together, these results and those obtained from localization of average nAPOGEE scores, suggest that T- and D-loop regions tend to accumulate variants with a similar nAPOGEE low score, whereas in the other tRNAs secondary structure motifs there is a wide variability in the nAPOGEE score distribution.

We then sought for the highest nAPOGEE scored regions in rRNAs. Therefore, we found that they were mostly located into two functional regions: in the peptidyl transferase center (PTC), mainly in two of its helices, i.e. the A-loop^48^, which is crucial for accommodation of the incoming aminoacyl-tRNA and, therefore, elongation of nascent protein^49^, and the H90 helix; in the H44 helix, a region containing two adenines that are universally conserved in all ribosomes and play a key role in mRNA translation^50^. We then applied the same autocorrelation analysis to rRNA variants, treating *MT-RNR1* and *MT-RNR2* as a single-RNA complex (**Fig. 3b**) and observed, as for tRNAs, positive spatial autocorrelation (permutation *q-value*<0.0001). Mapping the positive LISA values onto the complex highlighted functional regions in *MT-RNR2*, such as the PTC and the P-loop, and in *MT-RNR1*, such as the H44 helix, where stems showed a more deleterious score compared to loops (**Fig. 3b**). The results obtained via spatial autocorrelation are particularly compelling, as they highlight functional regions within rRNAs that remain detectable even when the spatial context is omitted. Overall, results obtained on rRNAs further demonstrate the presence of regions intolerant to variations, which can be especially found in regions binding tRNAs^36^.

**Extended Data Fig. 3.**
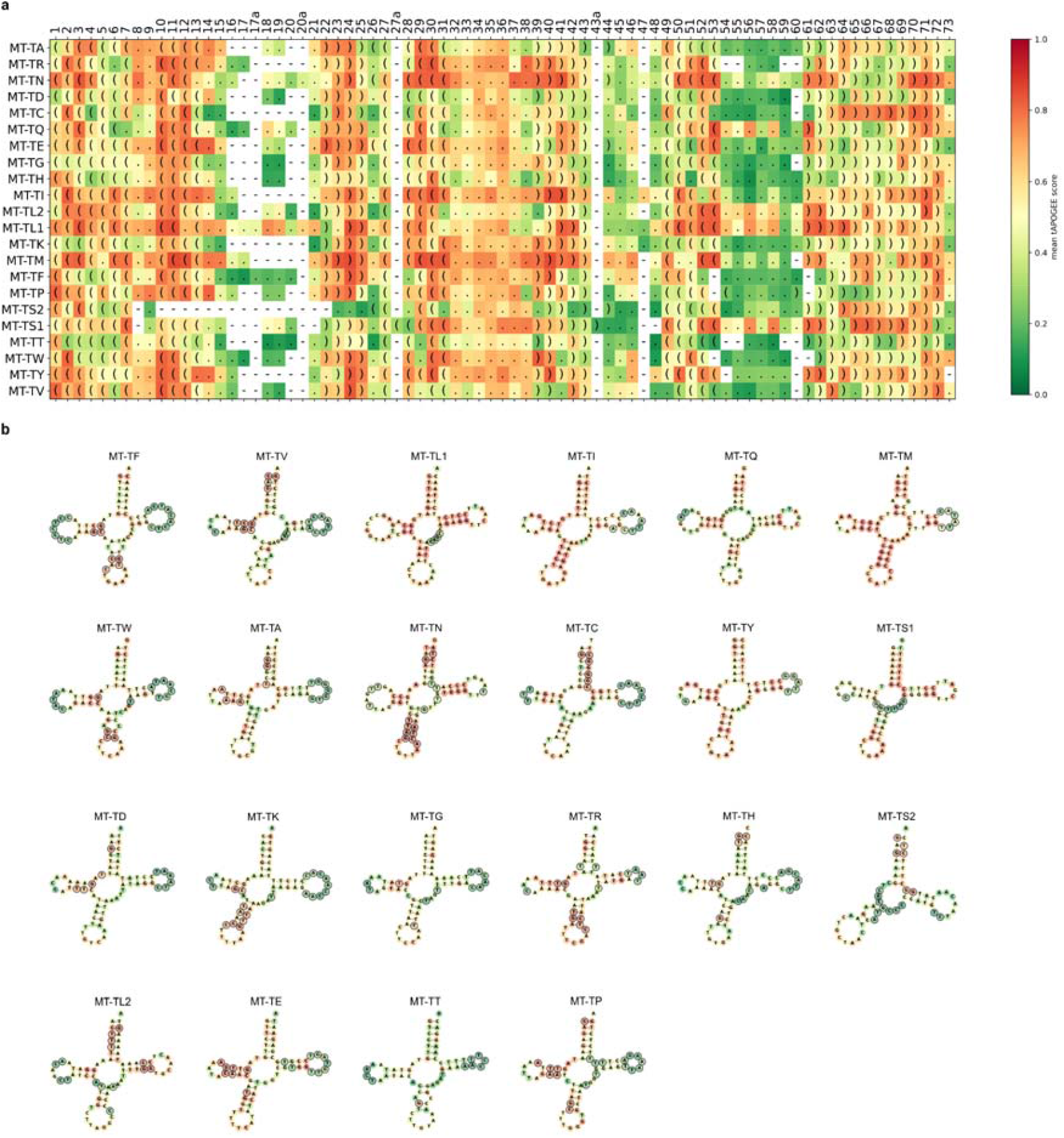
tRNAs nAPOGEE scores and spatial autocorrelation. **a**, nAPOGEE scores across tRNAs dot-bracket secondary structures, aligned according to the human MSA. **b**, nAPOGEE average scores for each residue across individual tRNAs secondary structures. nAPOGEE scores are represented in a color scale from green to red (i.e., from lower to higher scores). Residues showing significantly positive LISA values have been outlined in black.

### Probabilistic Interpretation of nAPOGEE Scores within ACMG/AMP Guidelines

To enhance clinical applicability and integration with existing variant interpretation frameworks, we converted raw nAPOGEE scores into posterior probabilities of pathogenicity, aligning the outputs with the computational evidence criteria recommended by the ACMG/AMP guidelines. This transformation was accomplished by empirically modeling the score distributions for benign and pathogenic variants separately. Specifically, we used ClinGen benign variants (**Dataset 4**) to derive the likelihood distribution for benign and likely benign SNVs, whereas the pathogenic distribution was modeled using ClinGen pathogenic or likely pathogenic variants (**Dataset 3**) supplemented by MITOMAP-confirmed and reported disease-associated variants (**Dataset 1**). In instances of limited variant counts, we employed OOB predictions to minimize overfitting and ensure an unbiased estimation of these empirical score distributions.

The posterior probabilities of pathogenicity were subsequently calculated using Bayes’ theorem. These probabilities were then mapped onto qualitative interpretative categories consistent with ACMG/AMP standards, facilitating standardized and clinically meaningful interpretations of in silico variant pathogenicity assessments (**Fig. 4a,d**). Comparing posterior probabilities between tRNAs and rRNAs, we observed that they are generally lower for rRNAs compared to tRNAs (**Fig. 4a,d**). This discrepancy is likely due to the limited biological understanding and functional annotation of potentially pathogenic variants within the rRNA regions^23,36^.

**Fig. 4.**
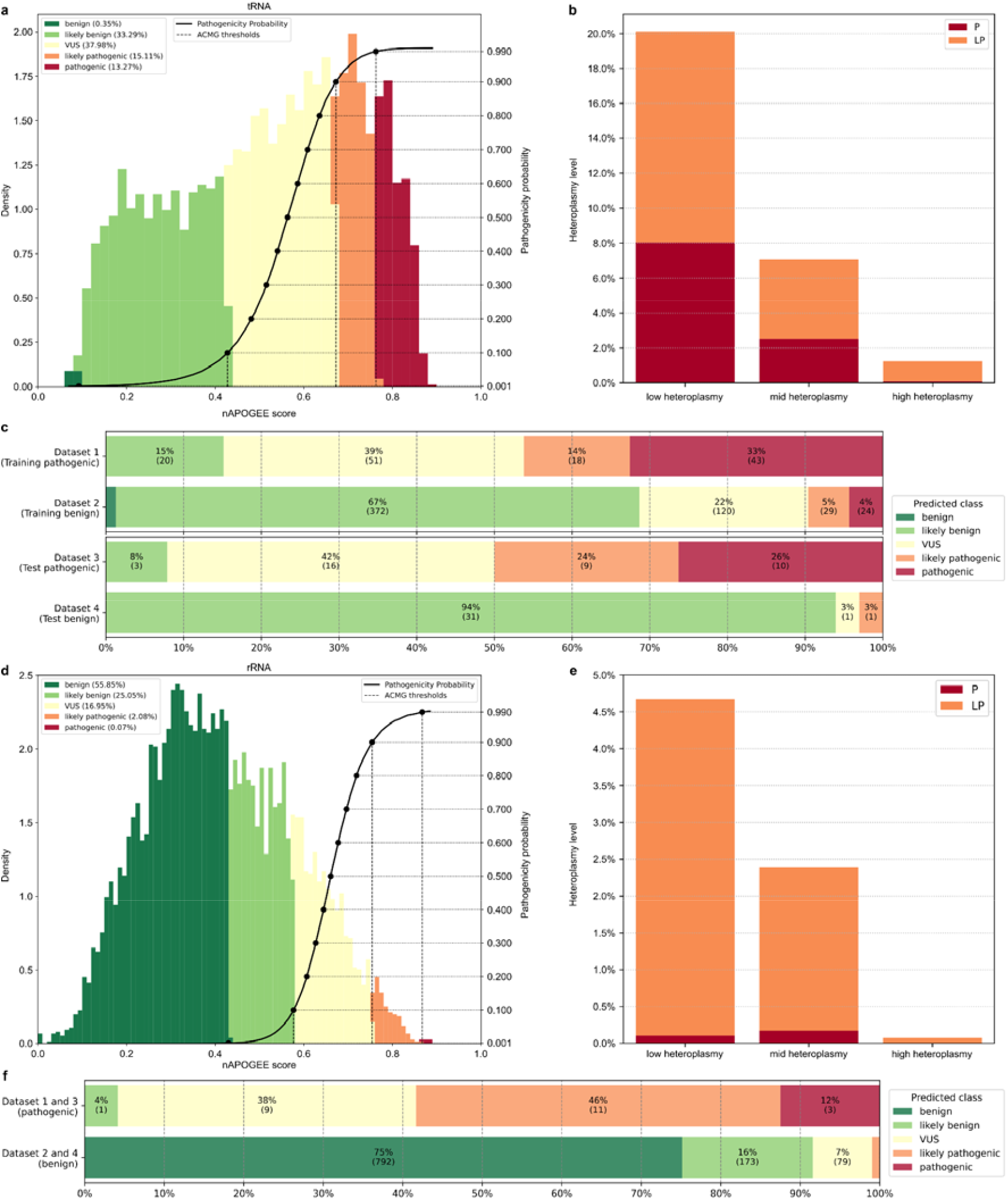
tRNA and rRNA variants classification according to ACMG/AMP guidelines. **a-c**, tRNA classification and **d-f**, rRNA variants classification according to ACMG/AMP guidelines. **a, d**, tRNA and rRNA variants classification using the posterior probabilities of pathogenicity. Classes were defined as previously done^22^: benign ≤0.001, 0.001 < likely benign ≤ 0.1, 0.9 ≤ likely pathogenic < 0.99, pathogenic ≥ 0.99. The remaining variants were defined as VUS (Variants of Uncertain Significance). **b, e**, Variants classification based on their level of heteroplasmy found in the gnomAD population. Levels of heteroplasmy were defined as following: low heteroplasmy variants (heteroplasmy ≤20%), middle heteroplasmy (>20% and ≤80%) and high heteroplasmy (>80%). **c, f**, tRNA and rRNA variants reclassification.

Focusing on tRNAs, our results showed that among all the possible SNVs, only a few were classified as benign (i.e., 16, 0.35%, **Fig. 4a**), while approximately 30% (i.e., 1283) of SNVs could be classified as likely pathogenic and pathogenic. These results are particularly noteworthy considering that, to date, ClinGen expert-curated tRNA SNVs include a comparable number of benign and pathogenic variants, with 33 and 38 SNVs reported in **Datasets 4** and **3**, respectively. Our analysis thus further supports the notion that tRNA genes are highly conserved^36^, and that sequence changes in these molecules can exert a significant biological impact. We further evaluated the misclassification rate of tRNA SNVs by analyzing the reclassification of SNVs in our datasets according to their unbiased pathogenicity probabilities (**Fig. 4c**). In particular, analyzing ClinGen-curated pathogenic dataset (**Dataset 3**), tAPOGEE misclassifies 8% of these variants (i.e., 3/38) into likely benign variants, while it correctly classifies 94% of ClinGen-curated benign variants (**Dataset 4**) into likely benign. These results underscore the inherent complexity in interpreting such variants.

Looking at rRNAs, we found that nAPOGEE posterior probabilities indicated that 162 (2.15%, **Fig. 4d**) of all possible SNVs could be classified as likely pathogenic or pathogenic variants. Since, to date, only two variants have been associated with clinical conditions (i.e., m.1555 A>G and m.1494 C>T), these results further reflect the necessity to develop dedicated tools to interpret rRNA variants that should be carefully investigated as they could exert a biological and functional role. Moreover, rAPOGEE classifies as benign and likely benign almost 81% of all possible SNVs (**Fig. 4d**). Results obtained from misclassification analysis, performed unifying **Dataset 1** and **3** and **Dataset 2** and **4** (**Fig. 4f**), due to the limited number of ClinGen-curated variants, further highlights that rAPOGEE correctly classifies the majority of benign variants (91%), suggesting that rRNA benign variants are easier to identify than pathogenic ones, likely due to our still limited biological understanding.

### Application to Population Data: gnomAD Heteroplasmy Stratification

To assess the distribution of potentially deleterious mitochondrial variants in the general population, we applied the ACMG/AMP-calibrated posterior probabilities derived from the nAPOGEE framework to all annotated noncoding SNVs present in the gnomAD mitochondrial dataset^51^. Variants were stratified according to their heteroplasmy levels to evaluate the relationship between predicted pathogenicity and population frequency (**Fig. 4b,e**). The analysis revealed a clear inverse relationship between the predicted pathogenicity and heteroplasmic fraction: variants classified as pathogenic or likely pathogenic were observed at progressively lower frequencies as heteroplasmy levels increased for both tRNAs (Kendall’s tau=-0.31, p-value<0.0001) and rRNAs (Kendall’s tau=-0.18, p-value<0.0001). This pattern is consistent with the action of purifying selection, which eliminates deleterious mitochondrial variants from the population, particularly at higher heteroplasmic fractions where the likelihood of phenotypic consequences is increased. These findings provide population-scale support for the biological plausibility of nAPOGEE predictions and reinforce the role of heteroplasmy as a key modulator of the pathogenicity of mitochondrial variants.

### Analysis of real-world tRNA and rRNA variants

We selected three Variants of Uncertain Significance (VUS) in tRNA and rRNA found in a local cohort of patients, showing clinical features consistent with a mitochondrial disorder, for which the nAPOGEE prediction provided an indication of pathogenicity or benignity, and analyzed them using molecular dynamics (MD) simulations. Specifically, the selected variants were m.611 G>A in *MT-TF* coding for tRNA-Phe, m.922 C>T in *MT-RNR1*, and m.3058 T>C in *MT-RNR2* coding for 12S and 16S rRNAs, respectively.

The first variant, m.611 G>A (*MT-TF*) was found at low heteroplasmic level (9%) in a muscle biopsy derived from a 62-year-old patient showing clinical symptoms of mitochondrial myopathy. The variant is not observed in individuals sequenced in the MITOMAP database. No other variants were identified in the mtDNA sequence, and no other genetic analyses were performed. The variant was classified as VUS, due to the low heteroplasmy found in the muscle, its undetectability in other tissues and in the patient’s relatives. By performing MD simulations of the m.611G>A tRNA-Phe mutant structure, we observed that the anticodon stem and loop (ASL) domain decreased in flexibility (**Fig. 5b**) compared to wild-type (WT) structure. This was mainly related to the high preservation of secondary structure integrity in the proximity of the anticodon (persistent base pairing between residues 28-33 and 40-45) compared to WT and a close likely benign variant, m.619 T>C (**Fig. 5c**). This behavior is reflected in their dynamic secondary structure, where m.611 G>A showed less flexibility than the WT throughout the simulation, with even the formation of pseudoknots that further reduced the entire domain flexibility (**Fig. 5d**). The MD result was consistent with the nAPOGEE prediction, which classifies the variant as likely pathogenic (nAPOGEE score 0.74, posterior probability 0.98). SHAP values of features contributing to this variant classification in the tAPOGEE pipeline were in agreement with MD results (**Fig. 5e**); indeed, feature contribution analysis underlined that the m.611 G>A stabilizes the tRNA secondary structure, while features as RNA-MSM and sequence conservation (highlighted by “Sequence Entropy” and “PhatsCons” and “PhyloP” scores) moved the variant toward a more deleterious effect.

**Fig. 5.**
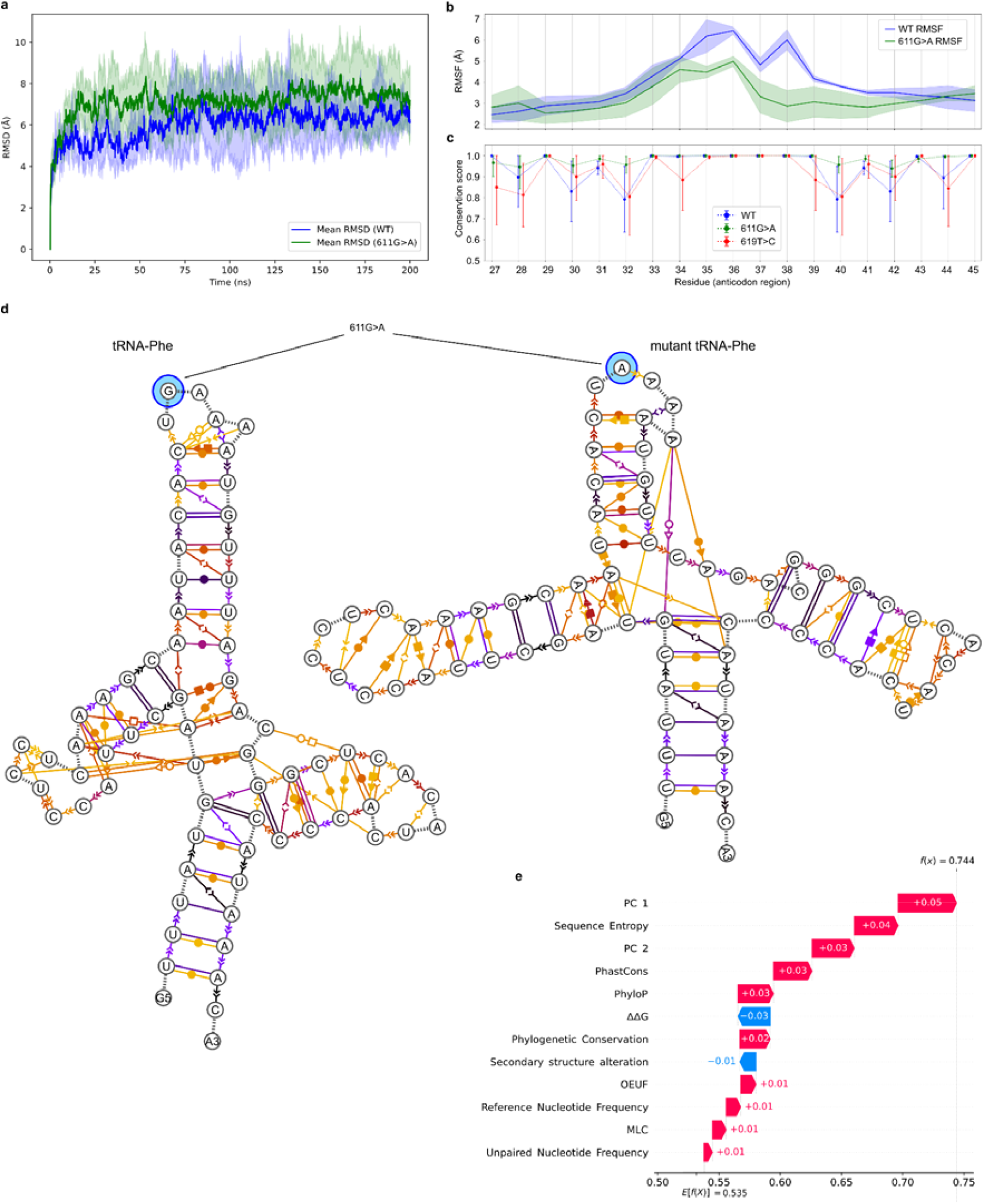
Structural dynamics and conservation of the mitochondrial tRNA-Phe anticodon region: analysis of m.611G>A. **a**, RMSD over 200 ns simulations for WT (blue) and m.611G>A (green); shaded areas represent the standard deviation across replicates. **b**, RMSF profiles of the anticodon region (residues 27–45). **c**, Anticodon secondary structure conservation during dynamics (mean ± 95% CI) for the WT tRNA, and those carrying the m.611 G>A variant and the likely benign m.619 T>C variant. **d**, Dynamic secondary structures of WT and m.611 G>A. Pairing occupancy is color-coded (yellow = 0%, black = 100%); geometric symbols indicate the base-pairing type and geometry (filled = cis, hollow = trans). **e**, SHAP values of the variant m.611 G>A contributing to its classification according to the tAPOGEE pipeline. Principal Components (PC) from 1 to 5 indicate the contribution of RNA-MSM, whose embedding channels underwent dimensionality reduction through a Principal Component Analysis (see Methods section). Features whose contributions are <0.01 were omitted; they are *PWM, PC 3-5, position in the human MSA, alternate nucleotide frequency, distance to PTM site, and coverage within MSA*. The overall nAPOGEE score assigned to the variant m.611 G>A was indicated in the figure as f(x) = 0.744.

We used the same analytical approach to estimate the preservation of the correct folding and tridimensional architecture also of rRNAs harboring two single nucleotide variants, m.922 C>T and m.3058 T>C. The m.922 C>T variant (in *MT-RNR1* gene coding for 12S rRNA) was identified in an adult patient presenting with ataxia, dysphagia, dysarthria, hearing impairment, peripheral neuropathy, and a positive family history (affected mother and sister). The variant was present at low heteroplasmic level in different biological samples (leukocytes 7%, fibroblasts 11%, bladder cells 5%). Whole Exome Sequencing (WES) was negative. The m.922 C>T variant has a frequency of 0.010% in MITOMAP and was classified as VUS. The m.3058 T>C variant (in *MT-RNR2* gene coding for 16S rRNA) was found in a child presenting with developmental delay, hypoglycemia, hepatomegaly, liver steatosis, after negative trio-WES. The variant was detected at 7% heteroplasmy in liver biopsy. The variant has a frequency of 0.002% in the MITOMAP database, is localized in a highly conserved nucleotide and was classified as VUS.

Our analyses disclosed that these variants affect the structural integrity of the 5’ domain of 12S rRNA (*MT-RNR1*) and the V domain of 16S rRNA (*MT-RNR2*). In 12S rRNA, the mutation replaces a non-canonical base pair between m.922 C and m.675 A with a canonical base pair (**Extended Data Fig. 4a**). This partially alters the conservation of the base pairing of the junction between helices H1 and H3 but does not visibly destabilize the 5’ domain compared to the WT and mutant simulations (**Extended Data Fig. 4b**). This was concordant with the nAPOGEE prediction (score 0.33, posterior probability 0.001) that suggested a likely benign classification for this variant. In contrast, the m.3058 T>C substitution in 16S rRNA introduced a non-canonical base pair in the arm of domain V, leading to a local decrease in structural integrity (**Extended Data Fig. 4c**). This was accompanied by reduced preservation of secondary structure elements in the loop region spanning nucleotides 1351-1370 and 1331-1350 (**Extended Data Fig. 4d**). nAPOGEE score indicates this variant as likely pathogenic (score 0.78, posterior probability 0.98), accordingly.

**Extended Data Fig. 4.**
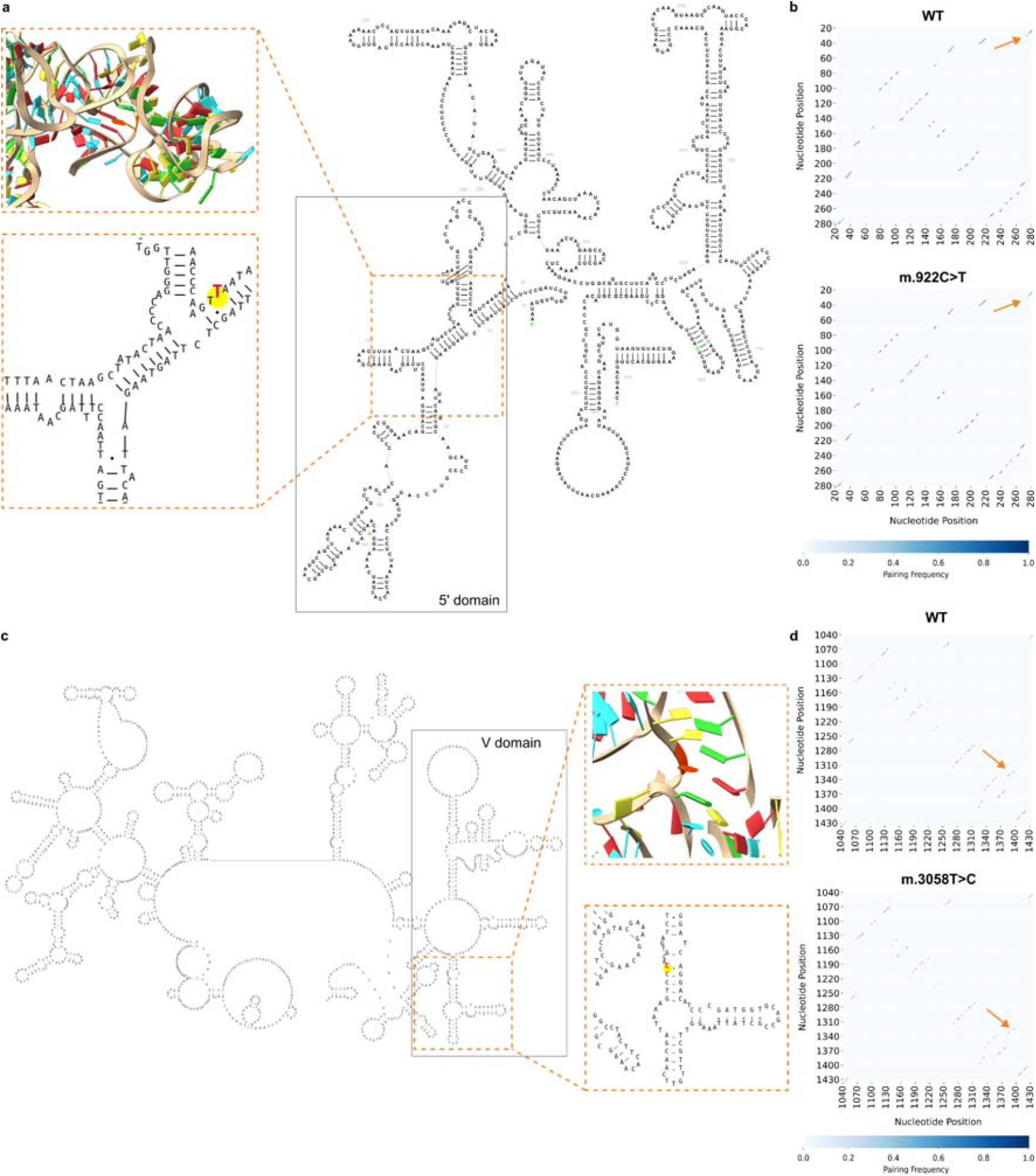
Structural and pairing analyses of WT and mutant mitochondrial 12S and 16S rRNAs. **a,** Schematic secondary structure of the mitochondrial 12S rRNA (*MT-RNR1), highligh*ting the 5′ domain and the local secondary and tertiary structures surrounding the nucleotide affected by the m.922 C>T mutation (in orange). **b**, Pairing frequency heatmaps derived from structural ensembles of 12S rRNA (*MT-RNR1) in the WT* (upper) and m.922 C>T mutant (lower). Arrows indicate the mutant region. **c**, Schematic representation of the secondary structure of the mitochondrial 16S rRNA (*MT-RNR2), highligh*ting the V domain and structural context of m.3058 T>C mutant (orange). **d**, Pairing frequency heatmaps for 16S rRNA (*MT-RNR2) in the WT* (upper) and m.3058 T>C mutant (lower). Arrows indicate the mutant region. The color intensity reflects the frequency of base pairing across conformational ensembles (scale: 0–1).

## Discussion

Accurate interpretation of mitochondrial noncoding variants remains a critical but underexplored challenge in genomic medicine. While the past decade has brought significant progress in computational tools for predicting the effects of missense variants in nuclear protein-coding genes, comparable advances in noncoding mitochondrial elements, particularly tRNAs and rRNAs, have lagged behind. Here, we present nAPOGEE, a unified framework for predicting the pathogenicity of all possible noncoding mitochondrial tRNA and rRNA variants. By combining evolutionary, structural, and embedding-based descriptors with Bayesian calibration aligned to ACMG/AMP guidelines^23^, nAPOGEE enables both mechanistic insight and clinically actionable classification, bridging a longstanding gap in variant interpretation.

Unlike their nuclear counterparts, mitochondrial tRNA and rRNA genes are present in single copies and lack functional redundancy, making them particularly vulnerable to the effects of deleterious mutations. Their central role in mitochondrial protein synthesis implies that even subtle sequence alterations can have pathogenic consequences. However, the systematic classification of such variants has been hampered by sparse datasets, absence of large-scale benchmarking, and lack of VEPs specifically designed for the mitochondrial genome. To date, there has been no specific VEP tailored to rRNAs. Conversely, for tRNA, following the ACMG/AMP recommendations^23^, we compared tAPOGEE with PON-mt-tRNA and MitoTIP, but not with HmtVar, since it was archived by its developers as of January 26, 2025, and is not currently being maintained. Our results demonstrate that tAPOGEE outperforms PON-mt-tRNA, even when evaluated on its original training dataset. However, when we compared tAPOGEE with MitoTIP, we observed that the latter showed better performance. This was largely expected, as MitoTIP integrates empirical data on variants, such as variant frequency in the general population and known pathogenicity status, which are now covered by mitochondrial ACMG/AMP guidelines (such as PM2, BA1, and BS1 for population frequencies, or PS3 and PM5 for clinical databases and literature evidence^23^). This advantage of MitoTIP in the classification of known mitochondrial variants makes it not generalizable to novel variants. Therefore, we compared tAPOGEE and MitoTIP using a less clinically characterized dataset. In this case, tAPOGEE achieved a better performance. Meanwhile, rAPOGEE fills a major void in the field as the first dedicated predictor for mitochondrial rRNA variants, enabling improved interpretation of variants implicated in disorders such as aminoglycoside-induced hearing loss (OMIM*561000), myopathy^52^, MELAS syndrome^53,54^, and hypertrophic cardiomyopathy^55,56^.

The strength of nAPOGEE lies not only in predictive accuracy but in its ability to uncover biological principles of mitochondrial genome function. Feature contribution analyses revealed that conservation drives pathogenicity in tRNAs, whereas spatial context dominates in rRNAs. Strikingly, RNA-MSM embeddings consistently contributed the strongest signal across both pipelines, even after collinear channels with annotated features were excluded. This indicates that latent structural and evolutionary constraints are implicitly encoded in the embedding space. Furthermore, Bayesian posterior calibration transforms raw model scores into ACMG/AMP-aligned probabilities, providing a direct path from algorithmic prediction to clinically meaningful classification.

Spatial autocorrelation analysis further revealed that pathogenicity signals are not randomly distributed but cluster within structurally and functionally constrained domains. In tRNAs, deleterious variants localize preferentially to stems whereas T-loops consistently accumulate neutral variants. In rRNAs, pathogenic clustering emerged in the peptidyl transferase center and helix 44, domains universally conserved across ribosomes. Importantly, these patterns persisted even after all explicit positional features were removed, demonstrating that pathogenicity can be inferred from correlated biological attributes without direct spatial annotation. This unified treatment allows the autocorrelation function to capture both intra- and inter-gene spatial dependence, reinforcing that structural domains constitute privileged targets of pathogenic variation. This observation aligns with the feature importance analysis, where localization-related features played a prominent role. Yet tAPOGEE’s broader feature space, integrating embeddings, intolerance scores, and structural descriptors, allowed the model to detect positional constraints implicitly, explaining its superior performance compared with predictors restricted to biochemical or conservation features.

A significant challenge in interpreting tRNA variants is to assess their effect on 3D structure, particularly on local structural flexibility. Although the 3D structures of RNA are primarily determined by their sequences, assessing their functional consequences remains complex, relying only on secondary or static tertiary structures. This makes molecular dynamics a valuable tool for investigating the conformational flexibility of RNA molecules. One particularly challenging region to interpret in tRNAs is the anticodon stem and loop (ASL) domain, where variants may exert their functional effect through, among others, alteration of the local structural flexibility, which can impair the fidelity and efficiency of translation. This is particularly relevant for the tRNA-Phe ASL domain functionality, where the impact of a variant of uncertain significance, m.611G>A, found in an in-house cohort, with symptoms of myopathy, was evaluated by analyzing its dynamical behavior, which disclosed the reduction in ASL domain flexibility. This effect may lead to multiple functional consequences including impaired codon-anticodon recognition, inefficient recognition by mitochondrial Elongation Factor Tu (EF-Tu), reduced aminoacylation, interference with post-transcriptional modifications, and increased susceptibility to degradation. This is consistent with some in vitro functional studies^57^, which demonstrated an approximately 100-fold reduction in the aminoacylation efficiency of tRNA-Phe by its tRNA synthetase (hmt-PheRS), while mitochondrial EF-Tu recognition was preserved, Phe codon decoding remained intact, and no evidence of A-to-I deamination was observed.

The m.611G>A variant, previously found in a patient with MERRF (Myoclonic Epilepsy with Ragged Red Fibers) syndrome, showing myopathy among other symptoms, was abundant in muscle (91%) but undetectable in all the other tissues tested and in familiars^58^. Currently, the variant is annotated as “reported” in MITOMAP and classified as “VUS” according to ClinGen experts. The nAPOGEE score classifies it as likely pathogenic, and MD simulations revealed a potential mechanism of pathogenicity, that is, local destabilization that may underlie the observed aminoacylation defects. Taken together, these results suggest that this variant could exert a biological role even at low heteroplasmic levels even if other functional assays are required. Notably, in vitro experiments^57^ showed that the migration profile of the mutant tRNA on native gels was comparable to that of the wild-type, suggesting a conserved global folding. This supports the idea that a mutation in a tRNA may leave a secondary or tertiary structure intact in static models but alter dynamic behavior and flexibility that can only be detected through MD simulations.

Similarly, structural evaluation is fundamental for describing the impact of mtDNA variants on the functioning of the human mito-ribosome^15,59^. We employed MD to estimate the folding and tridimensional architecture of rRNA harboring two single nucleotide variants with uncertain significance, m.922 C>T and m.3058 T>C, in 12S and 16S rRNAs, respectively, found in one of our cohorts. The m.922 C>T variant showed only a slight effect on the H1-H3 junction without compromising the global stability of the molecule, which is consistent with the nAPOGEE classification as likely benign. In contrast, the m.3058 T>C variant causes local destabilization and disruption of structural elements in loop regions, aligning with the nAPOGEE score, indicating pathogenicity. Although no functional studies are available for these cases, nAPOGEE allowed us to classify these variants of uncertain significance in two opposite directions, and MD supported this prediction by providing insight into the potential mechanisms of molecular pathogenesis. Functional validations and further investigations will be necessary to confirm this observation on this and other tRNA and rRNA variants.

Finally, the application of nAPOGEE to gnomAD population data revealed a striking inverse correlation between predicted pathogenicity and heteroplasmy levels, consistent with purifying selection against deleterious mitochondrial variants. This finding not only validates nAPOGEE predictions at population scale but also highlights heteroplasmy as a key modulator of mitochondrial disease risk. By aligning predictions with ACMG/AMP posterior probabilities, nAPOGEE offers a framework for systematic reclassification of variants of uncertain significance, supporting both diagnostic pipelines and future large-scale studies of mitochondrial genome variation.

Although nAPOGEE represents a significant advance, certain limitations must be acknowledged. The performance of rAPOGEE is constrained by the limited availability of well-characterized pathogenic rRNA variants. Although our cross-validation approach maximizes utility from the available data, prospective validation using emerging clinical cases is essential to confirm generalizability. Moreover, the use of RNA-MSM embeddings, while powerful, introduces dependencies on alignment quality and phylogenetic depth, areas that could benefit from further optimization and standardization.

However, nAPOGEE can already be used in the prediction of mitochondrial tRNA and rRNA somatic variants, which have been recently found to play a key role in cancer^17^. Moreover, the architecture is well-suited for adaptation to other noncoding mitochondrial elements, such as the displacement loop region (also known as control region) or regulatory motifs, and even to nuclear-encoded noncoding RNAs, including lncRNAs or snoRNAs, where annotation remains sparse and predictive models are nascent. Furthermore, the incorporation of longitudinal clinical data, tissue-specific expression profiles, and emerging single-cell mitochondrial sequencing data can further enhance the interpretive power of the model. Another exciting frontier lies in the integration of multimodal patient-specific data, including RNA-Seq, metabolomics, and imaging data, to move beyond static predictions toward dynamic, patient-tailored variant assessments. Such efforts would support the development of a robust precision medicine pipeline for mitochondrial disorders, where in silico prediction becomes a foundational component of diagnostic and therapeutic decision-making.

## Data Availability

The datasets supporting the conclusions of this article are included within the article and its Supplementary Table files. The MITOMAP reference files are available from [https://www.mitomap.org/MITOMAP/resources]. The ClinGen “pathogenic” and “likely pathogenic” revised mitochondrial variants are available at [https://www.mitomap.org/foswiki/bin/view/MITOMAP/MutationsRNACfrm] while the “benign” and “likely benign” are available at [https://www.mitomap.org/MITOMAP/ClinGenApproved]. The nAPOGEE probabilities/classes of pathogenicity can be freely downloaded from MitImpact [https://mitimpact.mcb2lab.org/] and are available for all the possible 12,060 tRNA and rRNA mitochondrial SNVs, as described in Supplementary Information. MSA derived from tRNAdb are available in GitHub [https://github.com/mazzalab/nAPOGEE], as the database is not currently available. rRNA sequences were downloaded from MIDORI2 [https://www.reference-midori.info/] and selected using information coming from NCBI Organelle [https://www.ncbi.nlm.nih.gov/datasets/organelle/]. rRNA MSA was obtained using ClustalOmega available at [https://www.ebi.ac.uk/jdispatcher/msa/clustalo]. rRNA models to analyze 3D localization of SNVs were generated from AlphaFold [https://alphafoldserver.com]. The RNA-MSM language model is available at [https://github.com/yikunpku/RNA-MSM]. tRNAscan-SE and R2DT used to generate tRNA and rRNA secondary structures can be found at [https://github.com/UCSC-LoweLab/tRNAscan-SE] and [https://rnacentral.org/r2dt], respectively. ViennaRNA employed to compute the change in ΔΔG associated with each SNV is available at [https://github.com/ViennaRNA/ViennaRNA]. MitoVisualize database is available at [https://mitovisualize.org/]. PON-mt-tRNA precomputed scores are available at [https://github.com/mazzalab/nAPOGEE], as the original database is no longer functioning. MITOTIP scores are available at [https://www.mitomap.org/downloads/mitotip_scores.txt]. GnomAD is freely accessible at [https://gnomad.broadinstitute.org].

## Code Availability

The tAPOGEE and rAPOGEE machine-learning workflows are available from https://github.com/mazzalab/nAPOGEE.

## Acknowledgements

We acknowledge the Italian Ministry of Health (RC2025). Additional non-financial support was provided to TM by NVIDIA Corporation, the Amber team, and the European High Performance Computing Joint Undertaking (EuroHPC JU) under the project “MITOLOGY: Multi-scale Simulation of the Mitochondrial Respiratory Chain Subunits” (EuroHPC Regular Access Call - EHPC-REG-2023R01-129 - PRACE EU Cal).

We gratefully acknowledge Dr. Stephan Bernhart (Interdisciplinary Center for Bioinformatics, Leipzig University, Leipzig, Germany) for his kind assistance with the MSA data originally reported in tRNAdb, following the discontinuation of the website. We acknowledge Dr. Eleonora Lamantea, responsible for the diagnostic mtDNA procedures at the Institute of Neurology “Carlo Besta.”

## Author contributions

T.M. and V.C. conceived the study. S.D.B., A.G., M.A., A.V., M.M., N.L., and A.N. performed computational genomic analyses. F.P. and T.B. performed MD analyses. S.D.B performed machine-learning analyses. T.M. and T.B. supervised computational analyses. L.C. and D.G. provided clinical data and performed wet-lab experiments. V.P. and V.C. supervised the genetic interpretation of results. M.T.L. and S.Z. provided MITOMAP data and analyses. A.U., A.G., and D.C.W. supervised the study. S.D.B., A.G., V.C., and T.M., wrote the paper, with the involvement of all authors.

## REFERENCES

1. Burgess, R. W. & Storkebaum, E. tRNA Dysregulation in Neurodevelopmental and Neurodegenerative Diseases. Annu Rev Cell Dev Biol 39, 223–252 (2023).

2. Schoenmakers, E. et al. Mutation in human selenocysteine transfer RNA selectively disrupts selenoprotein synthesis. J Clin Invest 126, 992–996 (2016).

3. Bösl, M. R., Takaku, K., Oshima, M., Nishimura, S. & Taketo, M. M. Early embryonic lethality caused by targeted disruption of the mouse selenocysteine tRNA gene (Trsp). Proc Natl Acad Sci U S A 94, 5531–5534 (1997).

4. Turvey, A. K., Horvath, G. A. & Cavalcanti, A. R. O. Aminoacyl-tRNA synthetases in human health and disease. Front Physiol 13, 1029218 (2022).

5. Orellana, E. A., Siegal, E. & Gregory, R. I. tRNA dysregulation and disease. Nat Rev Genet 23, 651–664 (2022).

6. Henderson, A. S., Warburton, D. & Atwood, K. C. Letter: Ribosomal DNA connectives between human acrocentric chromosomes. Nature 245, 95–97 (1973).

7. Kresoja-Rakic, J. & Santoro, R. Nucleolus and rRNA Gene Chromatin in Early Embryo Development. Trends Genet 35, 868–879 (2019).

8. Perry, R. P. et al. Evolution of the transcription unit of ribosomal RNA. Proc Natl Acad Sci U S A 65, 609–616 (1970).

9. Fan, W. et al. Widespread genetic heterogeneity of human ribosomal RNA genes. RNA 28, 478–492 (2022).

10. Farley-Barnes, K. I., Ogawa, L. M. & Baserga, S. J. Ribosomopathies: Old Concepts, New Controversies. Trends Genet 35, 754–767 (2019).

11. Taanman, J. W. The mitochondrial genome: structure, transcription, translation and replication. Biochim Biophys Acta 1410, 103–123 (1999).

12. Rackham, O. & Filipovska, A. Organization and expression of the mammalian mitochondrial genome. Nat Rev Genet 23, 606–623 (2022).

13. Wen, H. et al. Mitochondrial diseases: from molecular mechanisms to therapeutic advances. Signal Transduct Target Ther 10, 9 (2025).

14. Kogelnik, A. M., Lott, M. T., Brown, M. D., Navathe, S. B. & Wallace, D. C. MITOMAP: a human mitochondrial genome database. Nucleic Acids Res 24, 177–179 (1996).

15. Elson, J. L. et al. The presence of highly disruptive 16S rRNA mutations in clinical samples indicates a wider role for mutations of the mitochondrial ribosome in human disease. Mitochondrion 25, 17–27 (2015).

16. Prezant, T. R. et al. Mitochondrial ribosomal RNA mutation associated with both antibiotic-induced and nonsyndromic deafness. Nat Genet 4, 289–294 (1993).

17. Zhao, H. et al. Maternally inherited aminoglycoside-induced and nonsyndromic deafness is associated with the novel C1494T mutation in the mitochondrial 12S rRNA gene in a large Chinese family. Am J Hum Genet 74, 139–152 (2004).

18. Boscenco, S. et al. Functionally dominant hotspot mutations of mitochondrial ribosomal RNA genes in cancer. Nat Genet 57, 2705–2714 (2025).

19. Castellana, S., Rónai, J. & Mazza, T. MitImpact: an exhaustive collection of pre-computed pathogenicity predictions of human mitochondrial non-synonymous variants. Hum Mutat 36, E2413–22 (2015).

20. Castellana, S. et al. MitImpact 3: modeling the residue interaction network of the Respiratory Chain subunits. Nucleic Acids Res 49, D1282–D1288 (2021).

21. Castellana, S. et al. High-confidence assessment of functional impact of human mitochondrial non-synonymous genome variations by APOGEE. PLoS Comput Biol 13, e1005628 (2017).

22. Bianco, S. D. et al. APOGEE 2: multi-layer machine-learning model for the interpretable prediction of mitochondrial missense variants. Nat Commun 14, 5058 (2023).

23. McCormick, E. M. et al. Specifications of the ACMG/AMP standards and guidelines for mitochondrial DNA variant interpretation. Hum Mutat 41, 2028–2057 (2020).

24. Jühling, F. et al. tRNAdb 2009: compilation of tRNA sequences and tRNA genes. Nucleic Acids Res 37, D159–62 (2009).

25. Leray, M., Knowlton, N. & Machida, R. J. MIDORI2: A collection of quality controlled, preformatted, and regularly updated reference databases for taxonomic assignment of eukaryotic mitochondrial sequences. Environmental DNA 4, 894–907 (2022).

26. Madeira, F. et al. The EMBL-EBI Job Dispatcher sequence analysis tools framework in 2024. Nucleic Acids Res 52, W521–W525 (2024).

27. Abramson, J. et al. Accurate structure prediction of biomolecular interactions with AlphaFold 3. Nature 630, 493–500 (2024).

28. Zhang, Y. et al. Multiple sequence alignment-based RNA language model and its application to structural inference. Nucleic Acids Res 52, e3–e3 (2023).

29. Lake, N. J., Zhou, L., Xu, J. & Lek, M. MitoVisualize: a resource for analysis of variants in human mitochondrial RNAs and DNA. Bioinformatics 38, 2967–2969 (2022).

30. Chan, P. P. & Lowe, T. M. tRNAscan-SE: Searching for tRNA Genes in Genomic Sequences. Methods Mol Biol 1962, 1–14 (2019).

31. Sweeney, B. A. et al. R2DT is a framework for predicting and visualising RNA secondary structure using templates. Nat Commun 12, 3494 (2021).

32. The RNAcentral Consortium et al. RNAcentral: a hub of information for non-coding RNA sequences. Nucleic Acids Res 47, D1250–D1251 (2018).

33. Lorenz, R. et al. ViennaRNA Package 2.0. Algorithms Mol Biol 6, 26 (2011).

34. Pollard, K. S., Hubisz, M. J., Rosenbloom, K. R. & Siepel, A. Detection of nonneutral substitution rates on mammalian phylogenies. Genome Res 20, 110–121 (2010).

35. Siepel, A. et al. Evolutionarily conserved elements in vertebrate, insect, worm, and yeast genomes. Genome Res 15, 1034–1050 (2005).

36. Lake, N. J. et al. Quantifying constraint in the human mitochondrial genome. Nature 635, 390–397 (2024).

37. Sprinzl, M., Horn, C., Brown, M., Ioudovitch, A. & Steinberg, S. Compilation of tRNA sequences and sequences of tRNA genes. Nucleic Acids Res 26, 148–153 (1998).

38. Bohnsack, M. T. & Sloan, K. E. The mitochondrial epitranscriptome: the roles of RNA modifications in mitochondrial translation and human disease. Cellular and Molecular Life Sciences 75, 241–260 (2017).

39. Lundberg, S. M. et al. Explainable machine-learning predictions for the prevention of hypoxaemia during surgery. Nature Biomedical Engineering 2, 749–760 (2018).

40. Niroula, A. & Vihinen, M. PON-mt-tRNA: a multifactorial probability-based method for classification of mitochondrial tRNA variations. Nucleic Acids Res 44, 2020–2027 (2016).

41. Sonney, S. et al. Predicting the pathogenicity of novel variants in mitochondrial tRNA with MitoTIP. PLoS Comput Biol 13, e1005867 (2017).

42. Yarham, J. W. et al. A comparative analysis approach to determining the pathogenicity of mitochondrial tRNA mutations. Human mutation 32, (2011).

43. Meng, E. C. et al. UCSF ChimeraX: Tools for structure building and analysis. Protein Sci 32, e4792 (2023).

44. Sarzynska, J., Popenda, M., Antczak, M. & Szachniuk, M. RNA tertiary structure prediction using RNAComposer in CASP15. Proteins 91, 1790–1799 (2023).

45. Case, D. A. et al. The Amber biomolecular simulation programs. Journal of Computational Chemistry 26, 1668–1688 (2005).

46. Lu, X.-J., Bussemaker, H. J. & Olson, W. K. DSSR: an integrated software tool for dissecting the spatial structure of RNA. Nucleic Acids Res 43, e142 (2015).

47. Bottaro, S. et al. Barnaba: software for analysis of nucleic acid structures and trajectories. RNA 25, 219–231 (2019).

48. Koripella, R. K., Sharma, M. R., Risteff, P., Keshavan, P. & Agrawal, R. K. Structural insights into unique features of the human mitochondrial ribosome recycling. Proc Natl Acad Sci U S A 116, 8283–8288 (2019).

49. Ott, M., Amunts, A. & Brown, A. Organization and Regulation of Mitochondrial Protein Synthesis. Annu Rev Biochem 85, 77–101 (2016).

50. Vila-Sanjurjo, A., Mallo, N., Atkins, J. F., Elson, J. L. & Smith, P. M. Our current understanding of the toxicity of altered mito-ribosomal fidelity during mitochondrial protein synthesis: What can it tell us about human disease? Front Physiol 14, 1082953 (2023).

51. Chen, S. et al. A genomic mutational constraint map using variation in 76,156 human genomes. Nature 625, 92–100 (2023).

52. Coulbault, L. et al. A novel mutation 3090 G>A of the mitochondrial 16S ribosomal RNA associated with myopathy. Biochem Biophys Res Commun 362, 601–605 (2007).

53. Hsieh, R. H., Li, J. Y., Pang, C. Y. & Wei, Y. H. A novel mutation in the mitochondrial 16S rRNA gene in a patient with MELAS syndrome, diabetes mellitus, hyperthyroidism and cardiomyopathy. J Biomed Sci 8, 328–335 (2001).

54. Li, J.-Y. et al. A follow-up study in a Taiwanese family with mitochondrial myopathy, encephalopathy, lactic acidosis and stroke-like episodes syndrome. J Formos Med Assoc 106, 528–536 (2007).

55. Prasad, G. N. et al. Novel mitochondrial DNA mutations in a rare variety of hypertrophic cardiomyopathy. Int J Cardiol 109, 432–433 (2006).

56. Liu, Z. et al. The novel mitochondrial 16S rRNA 2336T>C mutation is associated with hypertrophic cardiomyopathy. J Med Genet 51, 176–184 (2014).

57. Ling, J. et al. Pathogenic mechanism of a human mitochondrial tRNAPhe mutation associated with myoclonic epilepsy with ragged red fibers syndrome. Proceedings of the National Academy of Sciences of the United States of America 104, (2007).

58. Mancuso, M. et al. A novel mitochondrial tRNAPhe mutation causes MERRF syndrome. Neurology 62, (2004).

59. Vila-Sanjurjo, A. et al. Structural analysis of mitochondrial rRNA gene variants identified in patients with deafness. Front Physiol 14, 1163496 (2023).

60. Biagini, T. et al. Molecular dynamics recipes for genome research. Briefings in Bioinformatics 19, (2018).

61. Biagini, T. et al. Are Gaming-Enabled Graphic Processing Unit Cards Convenient for Molecular Dynamics Simulation? Evolutionary Bioinformatics 15, (2019).

62. Pasculli, B. et al. Hsa-miR-210-3p expression in breast cancer and its putative association with worse outcome in patients treated with Docetaxel. Scientific Reports 1, (2019).

