## Supplementary Table for "nAPOGEE: A machine-learning platform for clinically actionable pathogenicity assessment of all mitochondrial noncoding variants"

### Supplementary Information

### Supplementary notes

In this study, we developed nAPOGEE, a machine-learning tool to predict the pathogenicity of all possible 4524 single-nucleotide variants (SNVs) in mitochondrial tRNAs and all possible 7536 SNVs in mitochondrial rRNAs.

nAPOGEE predictions were carefully assessed for each tRNA and rRNA SNV as described below. nAPOGEE predictions were not specifically calculated for SNVs localized in two genomic coordinates:

- m.5826, which is an overlapping site between *MT-TY* (encoding for tRNA-Tyr) and *MT-TC* (encoding for tRNA-Cys), both coding on strand “-”. As reported by Reichert et al., 1998^1^, during tRNA-Tyr/tRNA-Cys precursor processing, tRNA-Cys retains the overlapping residue at its first position, while tRNA-Tyr's truncated 3′-end is completed through an editing reaction that adds the missing residue (i.e., A). Therefore, for the three SNVs potentially located in the last position of *MT-TY,* we did not provide nAPOGEE *ad-hoc* calculated predictions, as they could not be considered as mtDNA genetic variants and, therefore, outside the scope of this work. For these three SNVs, we provided the nAPOGEE scores calculated for the overlapping *MT-TC* SNVs.
- m.3107, whose reference sequence is indicated as a “N”. This nucleotide number was maintained through the release of the mitochondrial genome to avoid errors due to the correction of a duplicated base initially located in this genomic coordinate (<https://www.ncbi.nlm.nih.gov/nuccore/251831106>; <https://www.mitomap.org/MITOMAP/CambridgeReanalysis>). Therefore, for SNVs potentially located in this residue, we did not provide an nAPOGEE score because there was no annotated reference sequence.

### References

1. Reichert, A., Rothbauer, U. & Mörl, M. Processing and editing of overlapping tRNAs in human mitochondria. *The Journal of biological chemistry* **273**, 31977–31984 (1998).
